# ATP6AP2-to-MMP14, a key pathway for osteoblast to osteocyte transition

**DOI:** 10.1101/2022.04.27.489713

**Authors:** Lei Xiong, Hao-Han Guo, Jin-Xiu Pan, Xiao Ren, Daehoon Lee, Lin Mei, Wen-Cheng Xiong

## Abstract

Osteocytes, derived from osteoblast (OB)-lineage, occupy lacunae within bone matrix and exhibit unique morphology with dendrite-like projections to form an inter-connected network. Such a network is essential for osteocytes to monitor and orchestrate bone homeostasis. Thus, it is of considerable interest to investigate how osteocytes are formed and how they built the network. Here we provide evidence for ATP6AP2, an accessory subunit of V-ATPase, in OB-lineage cells to be critical for OB-to-osteocyte transition, and osteocyte distribution, maturation, and morphogenesis. Mice (ATP6AP2^Ocn-Cre^) that selectively deplete ATP6AP2 in OB-lineage cells results in altered osteocyte distribution and morphology, impaired osteocyte maturation and dendrite-like processes, increased osteocyte cell death and cortical woven bone formation. Further mechanistic studies identify MMP14 (matrix metalloproteinase-14) as a critical downstream of ATP6AP2 for osteocyte differentiation. ATP6AP2 interacts with MMP14 and promotes MMP14 surface distribution largely in immature- or osteoid-osteocytes, where this pathway regulates bone matrix remodeling and osteocyte differentiation. Expression of MMP14 into ATP6AP2 knock out OB-lineage cells in the mouse cortical bone could diminish the deficits in osteocyte maturation, dendrite-like process formation, and survival. These results thus demonstrate an un-recognized function of ATP6AP2 in promoting OB-to-osteocyte transition and uncover a pathway from ATP6AP2-to-MMP14 in osteocyte differentiation.

## Introduction

Osteocytes, the most abundant type of cells in mature bone tissue, are living inside hard, rock-like bone. They were considered as inactive cells for a long time. However, recent studies have demonstrated that osteocytes are densely interconnected, forming a neuronal dendrite-like network, active in protein synthesis and modification. Osteocytes play multiple important roles, including regulating osteoblast (OB)- and osteoclast (OC)-activity to control bone remodeling, promoting systemic phosphate and calcium homeostasis, sensing mechanical loading or unloading, and secreting factors or hormones for other organs such as heart, liver and kidney (Bonewald, 2011, 2017; Dallas, Prideaux, & Bonewald, 2013; Faul et al., 2011; Gorski et al., 2016; Pereira et al., 2009; Shen, Grimston, Civitelli, & Thomopoulos, 2015). Osteocytes are descended from mesenchymal stem cells through OB differentiation (Blair et al., 2017). Several different transitional stages during OB-to-osteocyte transition have been identified, which include pre-osteoblast, osteoblast, embedding osteoblast, osteoid-osteocyte, mineralizing osteocyte, and mature osteocyte (Dallas & Bonewald, 2010; Franz-Odendaal, Hall, & Witten, 2006). It was believed that embedding of OBs to become osteocytes is a passive process, and OBs are “buried alive” under the matrix produced by adjacent OBs (Franz-Odendaal et al., 2006; Nefussi, Sautier, Nicolas, & Forest, 1991). This view has been challenged by recent studies which have shown that the embedding mechanism may be an active and invasive process in which the formation of osteocyte lacuna and canaliculi requires matrix degradation (Holmbeck et al., 2005; Zhao, Byrne, Wang, & Krane, 2000). However, the precise molecular and cellular mechanisms by which an OB becomes embedded in the bone matrix to begin a new life as an osteocyte, and the molecular mechanisms that regulate osteocyte maturation and network building remain poorly understood.

ATP6AP2 (ATPase H^+^ transporting accessory protein 2), also called PRR (pro-renin receptor), plays important roles in multiple signaling pathways and in multiple organs (Arthur, Osborn, & Yiannikouris, 2021; Ichihara & Yatabe, 2019; Nguyen et al., 2002; Nguyen & Muller, 2010; Wu et al., 2016). It is initially identified as PRR, because it binds to pro-renin and renin, and induces the conversion of angiotensinogen to angiotensin I (Nguyen et al., 2002). Later, it is recognized as a critical regulator of Wnt/β-catenin signaling and V-ATPase (Buechling et al., 2010; Cruciat et al., 2010). ATP6AP2 interacts with Wnt receptors, LRP6 and frizzled, in HEK293 cells, *Xenopus laevis tadpoles*, and *Drosophila* (Buechling et al., 2010; Cruciat et al., 2010). Suppression of ATP6AP2 in HEK293 cells or in *Drosophila* reduces Wnt canonical (β-catenin) and non-canonical (PCP, planar cell polarity) signaling (Buechling et al., 2010; Cruciat et al., 2010). ATP6AP2 is also identified as an accessory subunit of V-ATPase, a proton pump containing macromolecular complex, regulating the V-ATPase protein complex assembly and activity (Beyenbach & Wieczorek, 2006; Kinouchi et al., 2010). In line with the view for multi-functions that ATP6AP2 plays, ATP6AP2 is implicated in the pathogenesis of numerous diseases, such as hypertension, pre-eclampsia, diabetic microangiopathy, acute kidney injury, cardiovascular disease, cancer, obesity, mental disorders (e.g., depression and post-traumatic stress disorder), and neurodegenerative diseases (e.g., Parkinson disease and Alzheimer disease) (Fukushima et al., 2014; Goldstein, Speth, & Trivedi, 2016; Greco et al., 2012; Ichihara et al., 2006; Ichihara & Yatabe, 2019; Kurlak, Mistry, Cindrova-Davies, Burton, & Broughton Pipkin, 2016; Oba-Yabana et al., 2018; Saigusa et al., 2015; Satofuka et al., 2009; Schafer, Peters, & von Bohlen Und Halbach, 2013; Stransky, Cotter, & Forgac, 2016; Tan et al., 2014; Valenzuela et al., 2010; Wang et al., 2017). However, its functions in bone or bone disorders remain elusive.

Here, we provide evidence that mice with selective depletion of ATP6AP2 in osteoblast (OB)-lineage cells (ATP6AP2^Ocn-Cre^) exhibit impaired osteocyte dendritic processes, increased immature- or osteoid-osteocytes, increased osteocyte cell death, and accumulation of cortical woven bone. The cortical woven bone phenotype is accompanied with increases in bone formation, clustered abnormal immature osteocytes and osteocyte precursors, but decrease in mature osteocytes, suggesting critical functions of ATP6AP2^Ocn-Cre^ in regulating osteocyte distribution and OB-to-osteocyte transformation. To understand the underlying mechanisms, we carried out proteomics analysis of ATP6AP2 KO OB-lineage cells (compared with that of control cells) and identified MMP14, whose cell surface distribution and activity are lost in ATP6AP2-KO OB-lineage cells in culture and in vivo. Expression of MMP14 into ATP6AP2-KO osteocytes not only restore MMP14 mediated collagen-degradative activity, but also attenuate the deficits in osteocyte distribution, dendritic processes, and maturation in ATP6AP2^Ocn-Cre^ mice. Further mechanistic studies suggest that ATP6AP2 interacts with MMP14 and promotes MMP14 surface targeting. Taken together, these results suggest a function of ATP6AP2 in regulating MMP14 in immature osteocytes, revealing ATP6AP2 as a novel regulator of MMP14, and uncovering an unrecognized pathway, ATP6AP2 to MMP14, in OB-to-osteocyte transformation.

## Results

### Increased cortical woven bone mass in ATP6AP2^Ocn-Cre^ mice

To investigate ATP6AP2’s function in bone, we generated OB-selective ATP6AP2 conditional knockout mice, ATP6AP2^Ocn-Cre^, by crossing floxed ATP6AP2 (ATP6AP2^flox/X^) mice with osteocalcin (Ocn)-Cre (Fig. S1A), which expresses Cre largely in OB-lineage cells, including bone marrow stromal cells (BMSCs), OBs, and osteocytes (Pan et al., 2018; Wu et al., 2016; L. Xiong et al., 2015; M. Zhang et al., 2002). As expected, ATP6AP2 protein levels were selectively reduced in the OB-lineage, but not the osteoclast (OC)-lineage, cells in ATP6AP2^Ocn-cre^ mice, as compared with those of littermate control mice (Ocn-Cre mice) (Fig. S1B-E). Microcomputer tomographic (μCT) analysis of femurs from ATP6AP2^Ocn-cre^ mice (3-MO) showed increased cortical bone volumes/total volumes (Cb, BV/TV), compared with that of control littermates (Ocn-Cre) (Fig. 1A, B). However, the cortical bone material density (TMD, tissue mineral density) was significantly decreased (Fig. 1C), imply that the mutant mice may have an unhealthy cortical bone. We thus further examined the cortical bone structures by use of a combination of bone histological staining analyses. H&E staining showed marked and un-evened increases in the cortical bone thickness as well as many abnormal empty lacunae in ATP6AP2^Ocn-cre^ mice (Fig. 1D-F), supporting the view for an unhealthy cortical bone. Safranin-O staining (which is often used for the detection of cartilages) exhibited elevated, but abnormally distributed Safranin-O^+^ signals or cartilages in the cortical bones of the mutant mice (Fig. 1G, H), implicating a deficit in OB-to-osteocyte transformation. Picro-sirius red staining of bone sections (which were imaged by both bright field and polarized light) displayed features of “woven” bone in the mutant mice-containing many disorganized and thicker collagen fibers (Fig. 1I, J, deep red color). In contrast, the collagen fiber bundles appeared to be fine and regularly arranged in parallel in control mice (3-MO) (Fig. 1I, J, see yellow, orange and green color), exhibiting features of lamellar bone.

**Fig 1.**
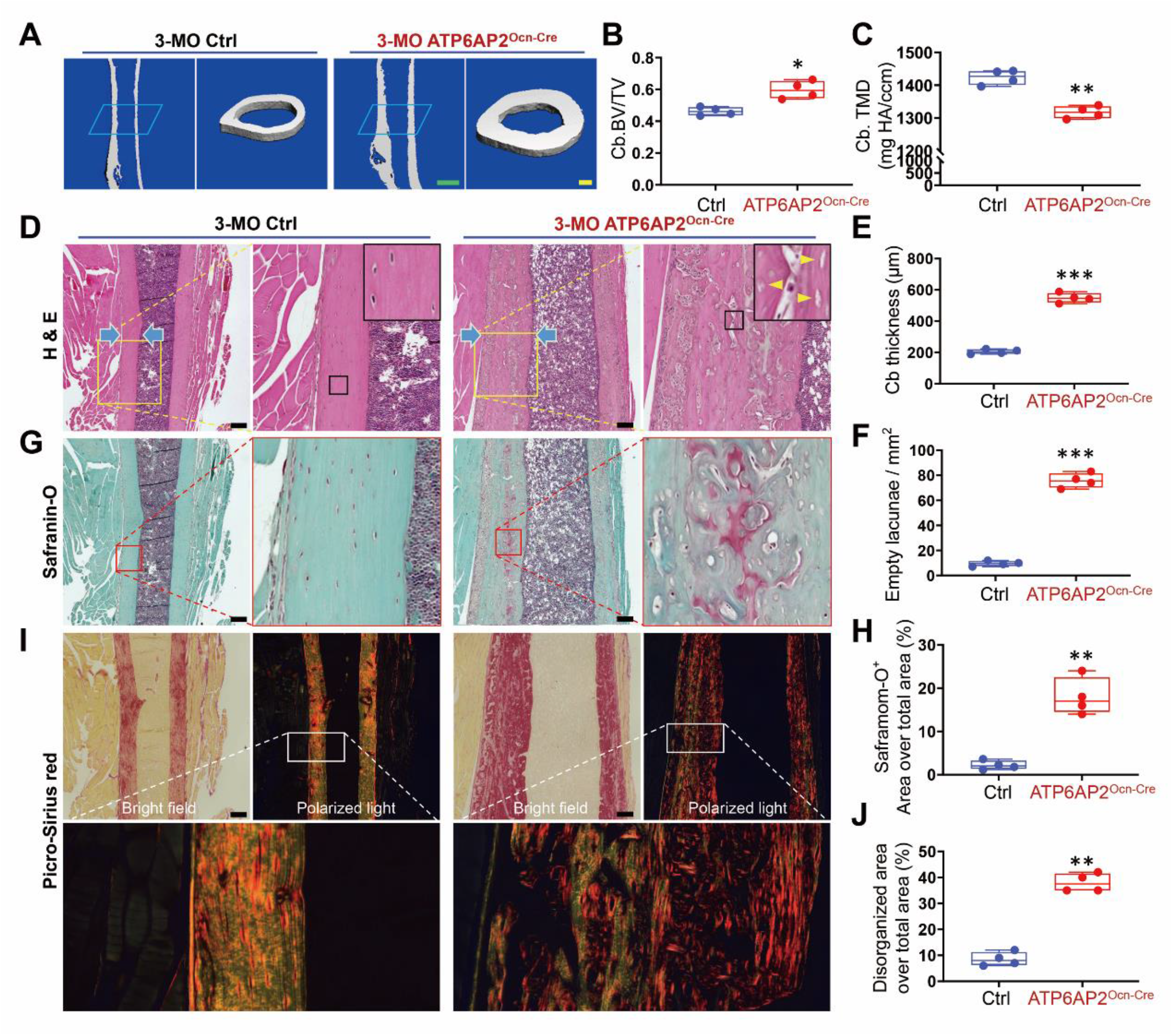
Increased cortical woven bone mass in ATP6AP2^Ocn-Cre^ mice. **(A)** Representative μCT 3D images of cortical bone from 3-MO ctrl and ATP6AP2^Ocn-Cre^ littermates. Green bar, 1 mm. Yellow bar, 200 μm **(B, C)** Quantification analyses of cortical bone (Cb) volumes over total volumes (BV/TV) and Cb material density (TMD, tissue mineral density) by direct model of μCT analysis. **(D)** Representative images of H&E stained cortical bone in the center region of femur sections from 3-MO ctrl and ATP6AP2^Ocn-Cre^ mice. Blue arrow indicted the cortical bone thickness; Yellow arrowhead indicted the empty lacunae. Bar, 150 μm **(E, F)** Quantification analyses of cortical bone thickness and empty lacunae. **(G)** Representative images of Safranin-O stained cortical bone from 3-MO ctrl and ATP6AP2^Ocn-Cre^ mice. Bar, 150 μm. **(H)** Quantification of cartilage distribution rate. **(I)** Representative images of Sirius red stained cortical bone from 3-MO ctrl and ATP6AP2^Ocn-Cre^ mice. Under polarized light, the collagen fiber bundles are fine and regularly arranged in parallel (yellow, orange and green color) in the ctrl mice, shows the features of lamellar bone. However, the collagen fibers appeared disorganized and thicker (deep red color) in the mutant mice, shows the features of woven bone. Bar, 150 μm. **(J)** Quantification of disorganized collagen distribution rate. Data in (B), (C), (E), (F), (H) and (J) are shown as box plots together with individual data points, and whiskers indicate minimum to maximum (n=4 animals per genotype). P values obtained by unpaired two-tailed t-test. *, P< 0.05. **, P< 0.01. ***, P< 0.001.

We further asked when these cortical bone phenotypes occurred in the mutant mice during postnatal development. H&E staining analysis showed obvious increases of cortical bone thickness and empty lacunae as early as P30 (1-MO) (Fig. 2A-C). In line with this result, Picro-sirius red staining showed that most cortical bones in the control mice developed lamellar bones at the age of 1-MO; however, in 1-MO ATP6AP2^Ocn-Cre^ mice, they contained a large number of woven bones (Fig. 2D, E), suggesting a role of ATP6AP2 in cortical bone development.

**Fig 2.**
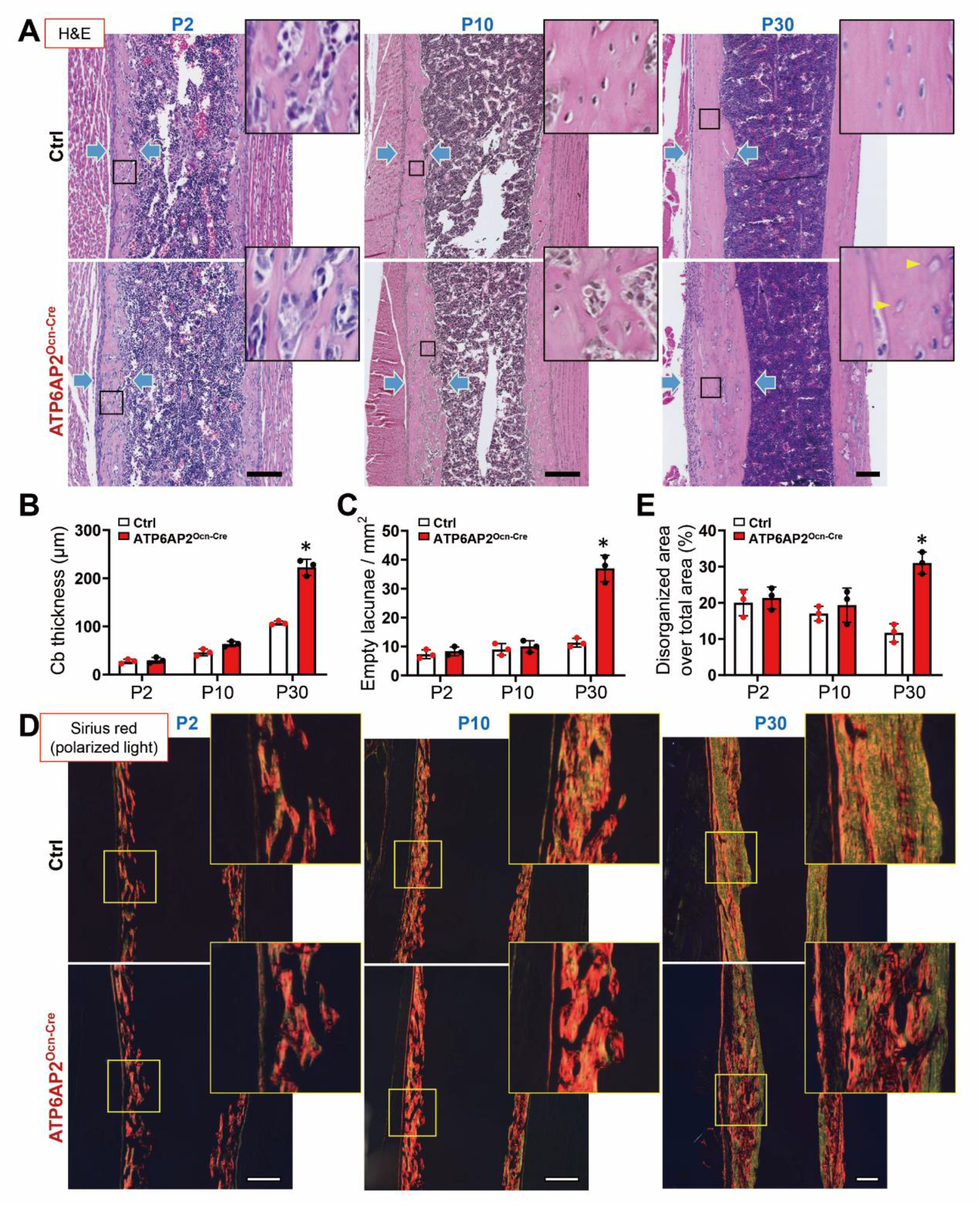
ATP6AP2^Ocn-Cre^ mice shown increased cortical bone thickness and empty lacunae from 1-MO. **(A)** Representative images of H&E stained cortical bone in the center region of femur sections from ctrl and ATP6AP2^Ocn-Cre^ mice at ages of P2, P10 and P30 (1-MO). Blue arrow indicted the cortical bone thickness; Yellow arrowhead indicted the empty lacunae. Bar, 150 μm **(B, C)** Quantification analyses of cortical bone thickness and empty lacunae. **(D)** Representative images of Sirius red stained cortical bone from ctrl and ATP6AP2^Ocn-Cre^ mice at ages of P2, P10 and P30 (1-MO). At 1-MO age, most cortical bone of the ctrl mice had formed lamellar bone, while the cortical bone of ATP6AP2^Ocn-Cre^ mice still had a large amount of woven bone. Bar, 150 μm. **(E)** Quantification of disorganized collagen distribution rate. Data in (B), (C) and (E) are presented as mean± SD (n=3 animals per genotype). *, P< 0.05 as determined by two-way ANOVA with Bonferroni post hoc analysis for multiple comparisons test.

### Elevated bone formation, accompanied with disorganized osteocyte distribution and increased cell proliferation and death, in ATP6AP2^Ocn-Cre^ cortical bone

To understand how ATP6AP2-KO in OB-lineage cells increases cortical woven bones, we examined bone formation in the cortical bones of the control and mutant mice by injecting fluorochrome-labeled calcein green into 1-MO mice twice at a 5-days interval. The mineral apposition rate (MAR), mineral surface/bone surface (MS/BS), and bone formation rate (BFR) in periosteum and endosteum bone regions were analyzed in non-decalcified femur bone sections from the injected mice. The MAR and BFR, but not MS/BS, were increased in both periosteum and endosteum cortical bone regions in ATP6AP2^Ocn-cre^ mice, as compared with those of control mice (Fig. S2A, B). Notice that there were calcein labeled mineralized areas in the center of the cortical bones in the ATP6AP2^Ocn-Cre^ mice, but not control mice (Fig. S2A, arrow heads), suggesting an abnormal cortical bone formation in the mutant mice.

We then crossed Ai9 mice, which express tdTomato in a Cre dependent manner, with Ocn-Cre and ATP6AP2^Ocn-cre^ mice to generate Ocn-Cre; Ai9 and ATP6AP2^Ocn-cre^; Ai9 mice, respectively, which allow us to visualize tdTomato^+^ bone cells in control (Ocn-Cre; Ai9) and mutant bones. In line with elevated cortical bone formation, the tdTomato^+^ (or Ocn-Cre^+^) cells were much more abundant in ATP6AP2^Ocn-cre^; Ai9 cortical bones (1-MO), as compared with those of littermate control mice (Ocn-Cre; Ai9) (Fig. 3A, B). Notice that the increased cortical bones were un-even, with a marked increase at the proximal diaphysis side in posterior face region (Fig. 3A, B). In addition, as shown in Fig. 3C and D, the tdTomato^+^ osteocytes were evenly distributed in the cortical bone of ctrl mice. However, in the mutant mice, these tdTomato^+^ cells were largely disorganized, many of the tdTomato^+^ cells were enlarged in their size, altered cell shape, clustered in their distribution, and localized largely in the region close to the periosteum (Fig. 3C-G), revealing sever disruptions in osteocyte distribution and morphology in ATP6AP2^Ocn-Cre^ mice.

**Fig 3.**
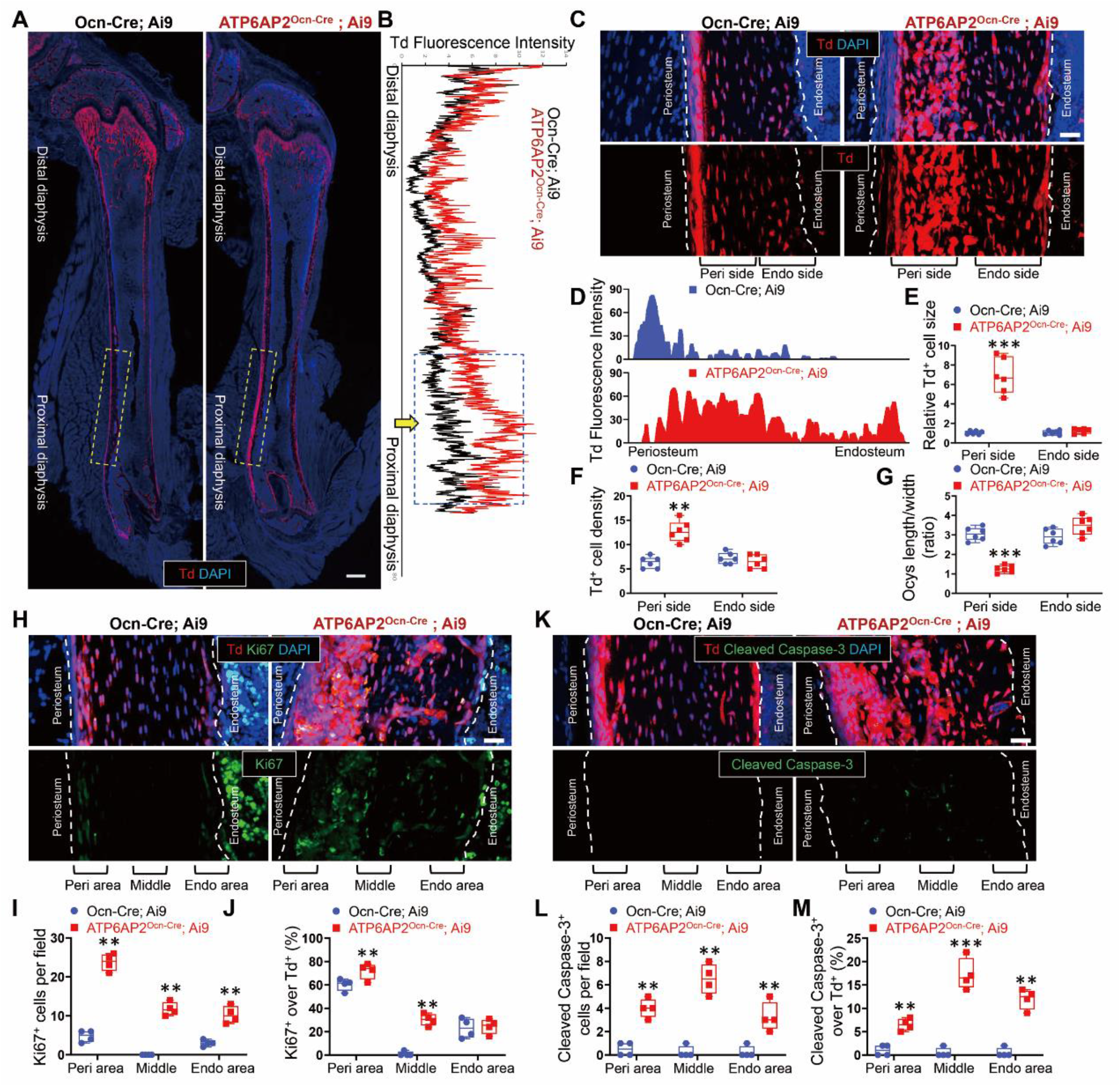
Disorganized osteocyte distribution and increased cell proliferation and death in the cortical bone of ATP6AP2^Ocn-Cre^ mice. **(A, B)** Td-Tomato expression on frozen femur sections from 1-MO Ocn-Cre; Ai9 and ATP6AP2^Ocn-Cre^; Ai9 mice. Bar, 300 μm. The Td-Tomato fluorescence intensity was shown in B. The dotted box indicated the marked increase of Td^+^ cells at the proximal diaphysis side in posterior face region. **(C)** Td-Tomato expression in the cortical bone region of femur sections from 1-MO Ocn-Cre; Ai9 and ATP6AP2^Ocn-Cre^; Ai9 mice. Bar, 50 μm. **(D-G)** Quantifications of the TdTomato red fluorescence intensity (D), the tdTomato^+^ cell size (E), cell density (F) and cell spherical shape (G). **(H-J)** Immunofluorescence of femur cortical bone in 1-MO Ocn-Cre; Ai9 and ATP6AP2^Ocn-Cre^; Ai9 mice by using Ki67 (cell proliferation marker) antibody. Bar, 20 μm. Quantification analysis was shown in I and J. **(K-M)** Immunofluorescence of femur cortical bone in 1-MO Ocn-Cre; Ai9 and ATP6AP2^Ocn-Cre^; Ai9 mice by using Cleaved caspase-3 (cell apoptosis marker) antibody. Bar, 20 μm. Quantification analysis was shown in L and M. Data in (E), (F), (G), (I), (J), (L) and (M) are shown as box plots together with individual data points, and whiskers indicate minimum to maximum (n=4 to 6 animals per genotype). P values obtained by unpaired two-tailed t-test. **, P< 0.01. ***, P< 0.001.

We further examined cell proliferation and cell death in ATP6AP2^Ocn-Cre^ cortical bones by immunofluorescence staining analyses. As shown in Fig. 3H-J, in the ctrl group, the Ki67^+^ proliferative cells were largely distributed in the bone marrow and the periosteum or endosteum area of the cortical bone, however, in the mutant mice, in addition to bone marrow, a dramatically increase in Ki67^+^ proliferative cells were detected in the cortical bone region (Fig. 3H, I). Note that more Ki67^+^ cells, co-labeled with tdTomato, were located in the periosteum side, implying that these enlarged cells in their size have more proliferative capacity (Fig. 3J). Additionally, the cleaved caspase-3^+^ apoptotic cells were also detected in the cortical bone of the mutant mice (Fig. 3K, L). Notice that more cleaved caspase-3^+^ cells, co-labeled with tdTomato, were distributed in the middle or on the periosteum side of the cortical bone, implying that cell death occurs mainly in small size osteocytes (Fig. 3M). These results suggest increased abnormal bone formation, cell proliferation and death in ATP6AP2^Ocn-Cre^ cortical bones, which may underlie the elevated cortical bone thickness and empty lacunae in the mutant mice.

### Impaired osteocyte maturation and dendrite-like processes in ATP6AP2^Ocn-Cre^ cortical bone

To better understand how the cortical bone phenotypes were developed in the mutant mice, it is necessary for further examine the osteocyte phenotype in detail. It is known that mature osteocytes (marked by SOST) are terminally differentiated cells derived from the OB-lineage precursors, which include Runx2 (Runt-related transcription factor 2)^+^ osteoblasts [also called osteoblastic osteocytes (Ocys)] and E11/Gp38^+^ immature Ocys (Bellido, 2014; Dallas et al., 2013) (Fig. 4A). To determine if and how ATP6AP2 regulates osteocyte differentiation, we examined SOST^+^ mature Ocys, E11/Gp38^+^ immature Ocys, and Runx2^+^ OBs/osteoblastic Ocys in the cortical regions of 1-MO Ctrl (Ocn-Cre; Ai9) and mutant (ATP6AP2^Ocn-Cre^; Ai9) mice. As shown in Fig. 4B, SOST^+^tdTomato^+^ mature Ocys were mainly distributed in the central region of the cortical bone in ctrl mice; and these SOST^+^tdTomato^+^ Ocys were marked reduced in the cortical region of ATP6AP2^Ocn-Cre^; Ai9 mice (Fig. 4B, C), indicating a decrease in mature Ocys. In contrast, the E11^+^tdTomato^+^ immature Ocys were increased and abnormally distributed in the mutant cortical bones (Fig. 4B, C). Whereas the E11^+^ immature Ocys were largely distributed in the region close to the periosteum of the control cortical bone, the E11^+^ Ocys were abnormally distributed in the center area of the cortical bone in the mutant mice (Fig. 4B). Further examination of Runx2^+^tdTomato^+^ OBs/osteoblastic Ocys also revealed a marked increase of these cells in the center area of the mutant cortical bones (Fig. 4B, C), with an altered distribution pattern (Fig. 4B). These results, a large number of OBs and immature osteocytes accumulated in the cortical bone of mutant mice, while the number of mature osteocytes was reduced (Fig. 4A), demonstrating a necessity of ATP6AP2 in OB-lineage cells for OB-to-osteocyte transition and differentiation.

**Fig 4.**
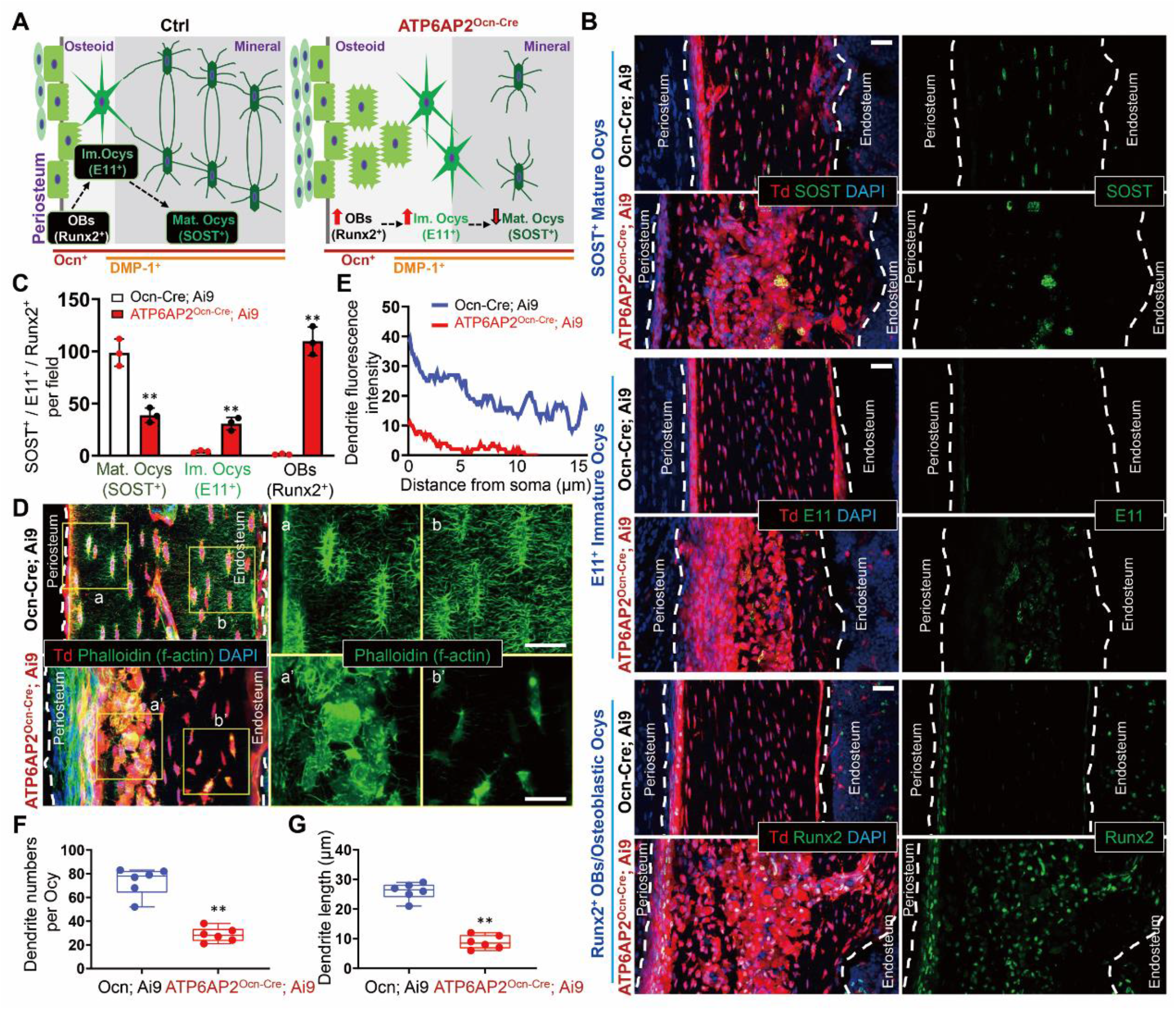
Impaired osteocyte maturation and dendrite-like processes in the cortical bone of 1-MO ATP6AP2^Ocn-Cre^ mice. **(A)** Schematic diagram depicting the transitional stages that occur as osteoblasts differentiate into mature osteocytes. The volume of the cell body and the number of cell organelles decreases during this process. **(B)** Immunofluorescence of femur cortical bone in 1-MO Ocn-Cre; Ai9 and ATP6AP2^Ocn-Cre^; Ai9 mice using SOST (mature osteocytes marker), E11 (also named gp38, early immature osteocyte marker) and Runx2 (preosteoblast, osteoblast, osteoblastic osteocyte marker) antibodies. Bar, 50 μm. **(C)** Quantifications of the percentage of SOST^+^, E11^+^ and Runx2^+^ cells per field. The values of mean ± SD (n = 3 animals per genotype) were shown. **, P< 0.01 as determined by unpaired two-tailed t-test. **(D)** Immunofluorescence staining of osteocytes and dendritic processes with phalloidin-488 for f-actin (green) in femur cortical bone of 1-MO mice with targeted deletion of ATP6AP2 in cells expressing osteocalcin (TdTomato^+^, ATP6AP2^Ocn-Cre^; Ai9) and their littermates (Ocn-Cre; Ai9). Right panel shows magnification of yellow boxed area of left panels. Bar, 20 μm. **(E-G)** Quantifications of the dendrite fluorescence intensity (E), the number of dendrites per osteocytes (F) and the osteocyte dendrite length (G). The values of mean ± SD (n = 3 animals per genotype) were shown in (C). Data in (F) and (G) are shown as box plots together with individual data points, and whiskers indicate minimum to maximum (n=6 animals per genotype). P values obtained by unpaired two-tailed t-test. **, P< 0.01.

We further asked whether the dendrite-like processes of the osteocytes, a morphological feature of mature osteoclasts, were affected. Using phalloidin staining to mark F-actin labeled dendrite-like processes (Dallas et al., 2013), we detected abundant well interconnected F-actin^+^ dendrite-like projections in the control (Ocn-Cre; Ai9, 1MO) cortical bone sections (Fig. 4D). These dendrite-like projections appear to be evenly distributed, without overlapping each other, exhibiting an organizational feature of “tiling”. However, in the mutant (ATP6AP2^Ocn-Cre^; Ai9, 1-MO) mice, little to no F-actin^+^ dendrite-like projections were detected in their osteocytes (Fig. 4D-G), indicating a sever disruption in osteocyte morphogenesis in ATP6AP2^Ocn-Cre^ mice, supporting the view for ATP6AP2’s function in promoting osteocyte maturation.

### Similar, but weaker, cortical bone deficit in ATP6AP2^DMP1-Cre^ mice

To further test the view for ATP6AP2 in the OB-lineage cells to regulate OB-to osteocyte transition and osteocyte differentiation, we generated ATP6AP2^DMP1-cre^ mice, by crossing ATP6AP2^flox/X^ mice with Dmp1-Cre, which is believed to express Cre largely in mature osteoblasts and osteocytes (Bellido, 2014; Dallas et al., 2013; Y. Lu et al., 2007; J. Xiong et al., 2015; J. Zhang & Link, 2016). Indeed, the td-Tomato^+^ fluorescence in DMP1-Cre; Ai9 cortical bone was largely detected in osteocytes (Fig. S3A); however, it was also detectable in muscles (Fig. S3A). Examining osteocyte distribution showed majority of tdTomato^+^ osteocytes to be evenly distributed in the cortical bone in ATP6AP2^Dmp1-cre^ mice (Fig. S3A, B), unlike that in ATP6AP2^Ocn-cre^ mice, which showed large number of tdTomato^+^ cells accumulated close to the periosteum side. However, small clusters of tdTomato^+^ cells were detectable in the mutant mice (Fig. S3A). These results suggest a little to weak deficit in osteocyte distribution when ATP6AP2 was depleted in Dmp1-Cre^+^ osteocytes. Further examining osteocyte phenotype showed similar, but weaker, osteocyte phenotypes, in ATP6AP2^Dmp1-cre^ mice to those in ATP6AP2^Ocn-Cre^ mice. For examples, the td-Tomato^+^ fluorescence intensity and cell density were higher in the cortical bone region in ATP6AP2^Dmp1-cre^ mice, as compared with those of control mice (Fig. S3A-C); the cortical bone thickness was increased in ATP6AP2^Dmp1-cre^ mice (Fig. S3D); and the reduced SOST^+^ mature osteocytes and increased E11^+^ and Runx2^+^ cells were noted in ATP6AP2^Dmp1-Cre^ mice, but less server than that in ATP6AP2^Ocn-Cre^ mice (Fig. S3E, F). Finally, imaging analysis of phalloidin/DAPI marked bone sections showed a decrease in the number of dendrites per osteocyte, but no significant change in dendrite length, in ATP6AP2^Dmp1-Cre^; Ai9 mice, as compared with those in control mice (Fig. S3G-J). These results suggest that ATP6AP2 in Dmp1-Cre^+^ mature osteoblasts and osteocytes is required for osteocyte maturation and the maintenance of the osteocyte network, but not osteocyte distribution, implicating a critical role of ATP6AP2 in Ocn-Cre^+^ OB/osteocyte precursors for osteocyte distribution, morphogenesis, and OB-to-osteocyte transition (Fig. 4A).

### Reduced MMP14 surface distribution and activity in ATP6AP2-KO OBs and immature osteocytes

To investigate molecular mechanisms underlying osteoblastic ATP6AP2 promoting OB-to-osteocyte transition, we carried out proteomic analysis by liquid chromatography-tandem mass spectrometry to screen for altered plasma membrane proteins in primary cultured ATP6AP2-KO Ocn-Cre^+^ OB-lineage cells (Fig. 5A). Both ctrl (Ocn-Cre) and ATP6AP2-KO OB-lineage cells were incubated with NHS-biotin for 45 min at 4 °C to label cell surface membrane proteins. The biotin labeled surface proteins were pulled down by streptavidin-agarose beads and subjected to proteomic analysis and western blot analysis. Among a total of 542 proteins identified, 27 proteins were down-regulated and 2 proteins was up-regulated in ATP6AP2-KO OBs, as compared with those of ctrl (Fig. 5B). Heat map analysis of these 27 down-regulated and 2-up-regulated proteins was illustrated in Fig. 5C. Gene ontology (GO) molecular functional analysis of these dysregulated proteins exhibited different biological processes, including cell adhesion molecules, cytokine binding proteins, and metallopeptidase activity (Fig. 5D). We further investigated MMP14, also called MT1-MMP, a protein belonging to the matrix metalloproteinase (MMP) family that is critical for the breakdown of extracellular matrix (ECM) (de Vos, Wong, Welting, Coull, & van Steensel, 2019; C. Lu, Li, Hu, Rowe, & Weiss, 2010). MMP14 was indeed markedly reduced in the plasma membrane fractions in ATP6AP2-KO OBs, as compared with that of controls, by Western blot analysis of biochemical cell fractions of OBs (Fig. 5E, F). This effect appeared to be selective, as the cell surface Lrp1 level was un-changed in ATP6AP2-KO cells (Fig. 5E, F). Notice that the total protein levels of MMP14 was also reduced in ATP6AP2-KO cells (Fig. 5E, G), while its mRNA level had little to no change (Fig. 5H), suggesting that ATP6AP2 may play a role in regulating MMP14 protein trafficking and stability. This view was further supported by the observations of lower cell surface levels of exogenous expressed MMP14-eGFP in ATP6AP2-KD (knock-down) MC3T3 cells than those of control MC3T3 cells (Fig. 5I, J). As MMP14 surface distribution is crucial for its activation, we further tested this view by examining MMP14 mediated degradation of the extracellular matrix (ECM)(e.g., type I collagen) in control (Ocn-Cre; Ai9) and mutant (ATP6AP2^Ocn-Cre^; Ai9) mice. As expected, the degradation of the fluorescent conjugated type I collagen was much lower in osteocytes derived from the mutant mice than those of control mice (Fig. 5K, L), indicating a reduced MMP14 activity in ATP6AP2-KO osteocytes.

**Fig 5.**
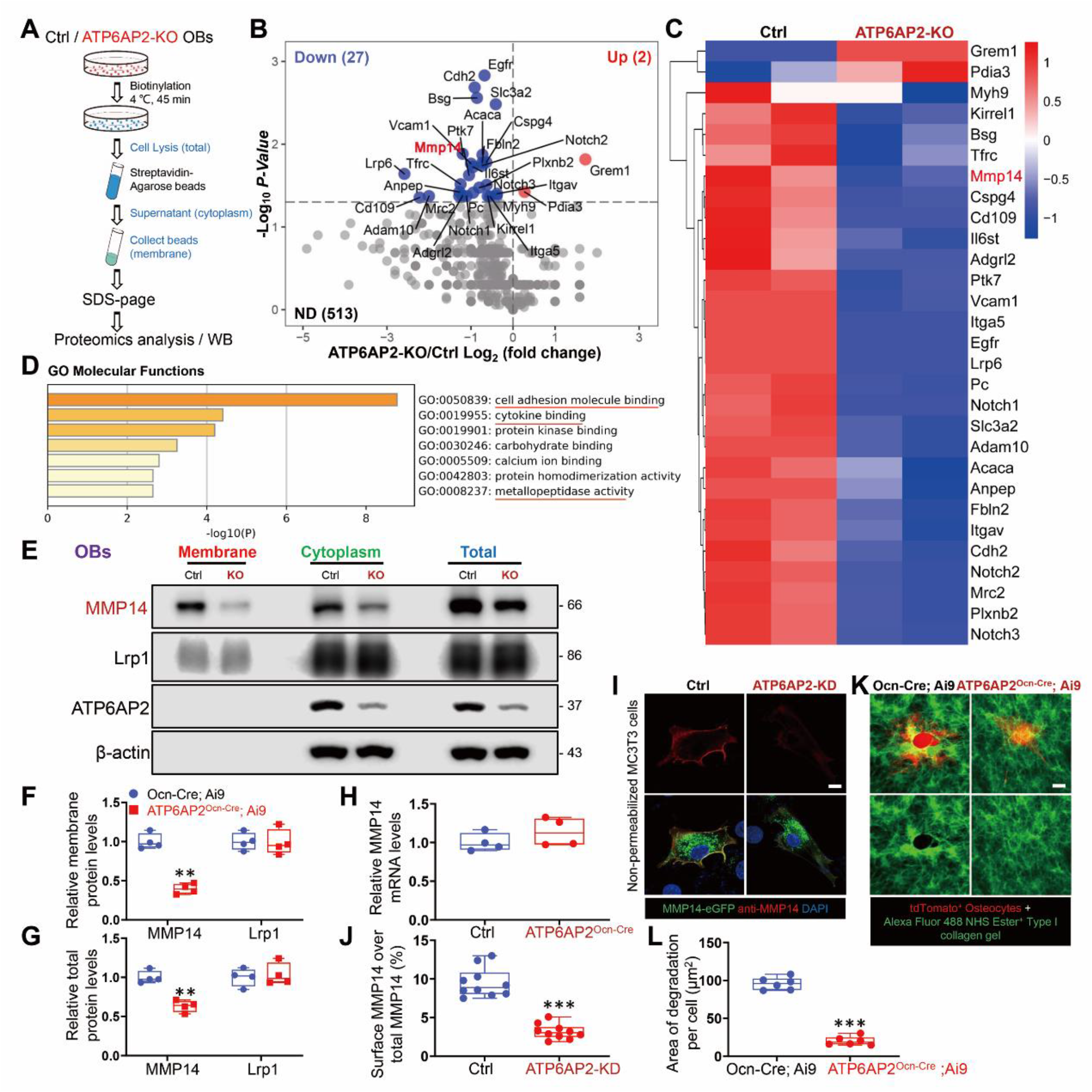
Reduced MMP14 surface distribution and activity in ATP6AP2-KO osteoblasts and immature osteocytes. **(A)** Schematic diagram of quantitative proteomic analysis. Proteomic data were obtained from surface proteins of ctrl and ATP6AP2-KO OBs. **(B)** Volcano plots of differentially expressed proteins in the cell surface of ctrl and ATP6AP2-KO OBs. The up-regulated proteins were marked in red, and the down-regulated proteins were indicated in blue (P < 0.05). **(C)** Heat map protein expression z-scores computed for the 29 proteins that are differentially expressed in cell membrane of Ctrl and ATP6AP2-KO OBs (P < 0.05). **(D)** The gene ontology (GO) molecular functional analysis shows the top 7 significant enrichment terms represented by molecular function, with longer column representing more significant enrichment. MMP14 was involved in many biological processes, including cell adhesion molecule binding, cytokine binding and metallopeptidase activity. **(E-G)** Western blot analyses of lysates of total, cytoplasm and cell surface fractions of ctrl and ATP6AP2-KO OBs. Quantification of the data was shown in F and G. **(H)** Real-time PCR analysis of MMP14 expression in BMSCs of 3-MO ctrl and ATP6AP2^Ocn-Cre^ mice. **(I, J)** Immunostaining analysis of cell surface MMP14 in non-permeabilized ctrl and ATP6AP2-KD MC3T3 cells transfected with the MMP14-eGFP plasmid. Bar, 10 µm. Quantification analysis was shown in J, the ratio of cell surface MMP14 over total MMP14. **(K, L)** Representative images (K) and quantitative measurement (L) of type I collagen layer degradation assay of osteocytes derived from Ai9; Ocn-Cre and Ai9; ATP6AP2^Ocn-Cre^ mice. The type I collagen gel was labeled by Alexa fluor 488 NHS ester dye before seeding the cells. Confocal micrographs showing degradation of the fluorescent collagen layer, indicated by a dark hole, in ctrl osteocytes, but not in ATP6AP2-KO osteocytes. Bar, 10 μm. Data in (F), (G), (H), (J) and (L) are shown as box plots together with individual data points, and whiskers indicate minimum to maximum (n=4 or 6 animals per genotype in F, G, H and L, n = 10 cells in J). P values obtained by unpaired two-tailed t-test. **, P< 0.01. ***, P< 0.001.

We next examined whether in vivo MMP14 expression and distribution in osteocytes of cortical bones were affected by loss of ATP6AP2. Notice that MMP14 was abundantly expressed in osteocytes that are largely distributed in the periosteum region of the control mice (Ai9; Ocn-Cre) (Fig. 6A, B), implying MMP14 to be selectively expressed or enriched in immature or osteoid osteocytes or osteoblastic osteocytes. Indeed, co-immunofluorescence staining analysis showed that MMP14^+^ osteocytes were E11^+^, a marker for immature osteocytes (Fig. 6G, H). In contrast from the control mice, MMP14 in ATP6AP2^Ocn-Cre^; Ai9 mice was marked decreased and abnormally distributed, as those of E11^+^ immature osteocytes in the mutant mice (Fig. 6G, H). MMP14 was reduced in E11^+^ immature osteocytes in the mutant cortical bones (Fig. 6G, H). Higher power imaging analysis showed a vesicle-like distribution of MMP14 in the mutant osteocytes, instead of an even cell surface distribution in the ctrl osteocytes (Fig. 6D-F). These results thus provide in vivo evidence for ATP6AP2 to regulate MMP14 protein trafficking and surface targeting, likely in large in immature osteocytes.

**Fig 6.**
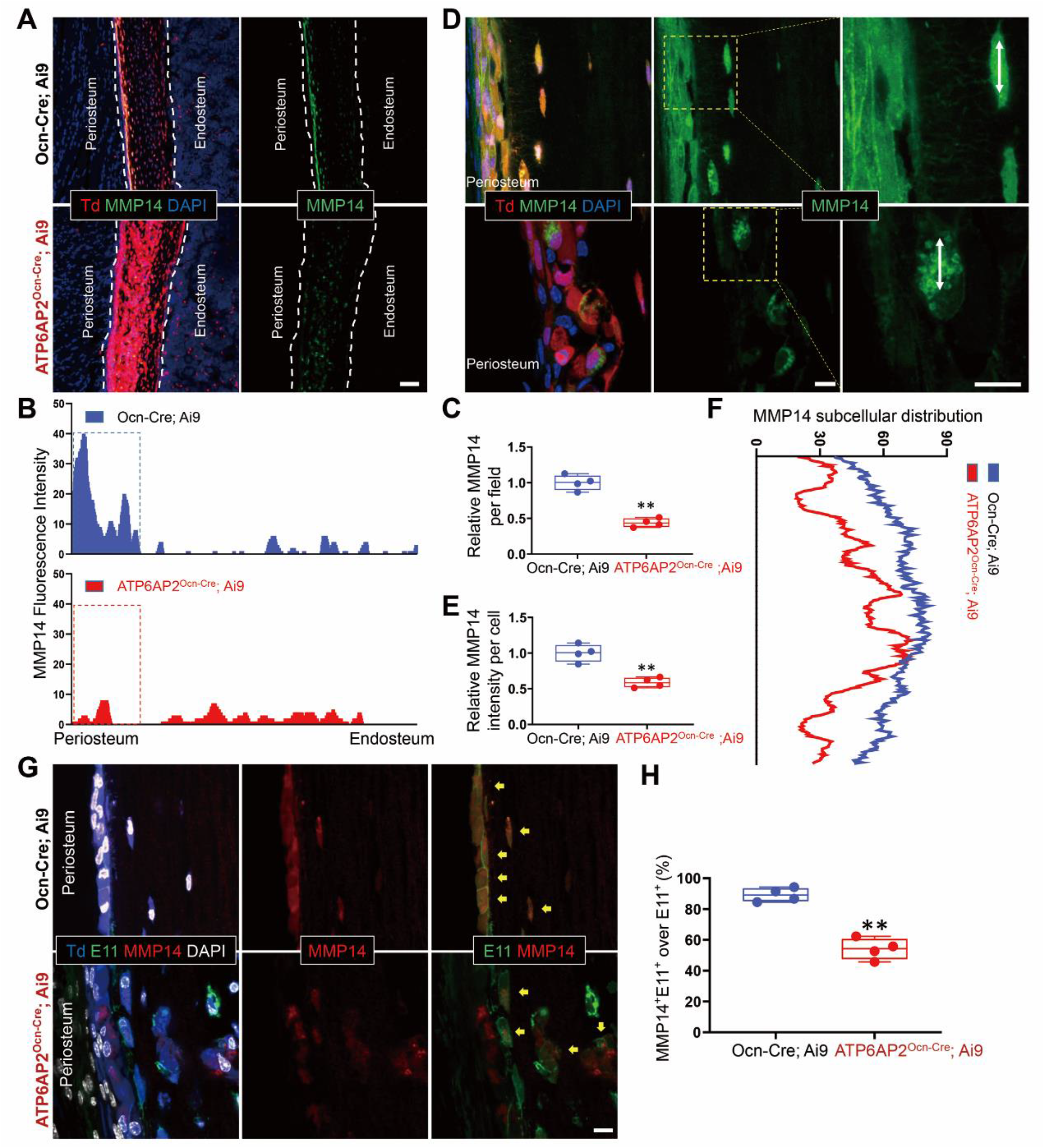
Reduced MMP14 surface distribution in ATP6AP2-KO osteocytes. **(A-C)** Immunofluorescence of femur cortical bone in 1-MO Ocn-Cre; Ai9 and ATP6AP2^Ocn-Cre^; Ai9 mice using MMP14 antibody. Representative images were shown in A. Bar, 50 μm. Quantification analyses of MMP14 fluorescence intensity was shown in B and C. **(D-F)** The vesicle-like distribution of MMP14 in ATP6AP2-KO cells. Immunostaining images of MMP14 in the periosteum area of 1-MO Ocn-Cre; Ai9 and ATP6AP2^Ocn-Cre^; Ai9 mice were shown in D. Bar, 20 μm. Quantification analyses of MMP14 fluorescence intensity per cells and fluorescence distribution in the marked cells were shown in E and F. **(G, H)** MMP14 high expressed in immature osteocytes. Immunofluorescence images of femur cortical bone in 1-MO Ocn-Cre; Ai9 and ATP6AP2^Ocn-Cre^; Ai9 mice using MMP14 and E11(gp38) antibodies were shown in G. Bar, 20 µm. Quantification analysis of the ratio of MMP14^+^; E11^+^ over total E11^+^ cells were shown in H. Data in (C), (E) and (H) are shown as box plots together with individual data points, and whiskers indicate minimum to maximum (n = 4 animals per genotype). **, P< 0.01 as determined by unpaired two-tailed t-test.

### MMP14 attenuation of the cortical osteocyte deficits in ATP6AP2^Ocn-Cre^ mice

To determine whether the reduced MMP14 is responsible for the cortical bone deficit in ATP6AP2^Ocn-Cre^ mice, it is necessary to address if expression of MMP14 into the mutant mice is capable to rescue the deficits. We thus generated lentivirus expressing MMP14-eGFP fusion protein in a Cre-dependent manner (Fig. S4A, B). Primary cultured OBs from ATP6AP2^Ocn-Cre^ mice infected with the lentivirus-MMP14-eGFP showed MMP14-eGFP fusion protein expression (Fig. S4C). As MMP14 surface targeting is crucial for its activation, we further tested this view by examining MMP14 mediated degradation of the extracellular matrix (ECM) (e.g., type I collagen). Control and ATP6AP2-KO OBs infected with or without lentivirus-MMP14-eGFP were cultured with fluorescence conjugated type I collagen gel. Whereas ATP6AP2-KO OBs showed a decrease in the collagen degradation, as compared with that of control OBs, infection of the lentivirus-MMP14-eGFP in the mutant OBs could diminish the deficit (Fig. S4D, E), supporting the view for ATP6AP2-MMP14 pathway in regulating collagen matrix degradation in culture.

We then asked whether injecting the lentivirus-MMP14-eGFP into ATP6AP2^Ocn-Cre^ cortical bones could diminish their bone deficits. The control and MMP14-eGFP lentiviruses were injected into the femur cortical bones of control and ATP6AP2^Ocn-Cre;^ Ai9 mice at age of P15 (Fig. S5A; Fig. 7A). One month later, the cortical bones were isolated and subjected to the co-immunostaining analysis. As shown in Fig. S6A and B, in the MMP14-EGFP virus injection side, GFP positive signals were detected in the tdTomato^+^ osteocytes, demonstrating the MMP14-eGFP expression. While no significant changes were detected on control osteocytes expressing MMP14-eGFP fusion protein (Fig. S5B-J), expressing MMP14-eGFP into the ATP6AP2^Ocn-Cre^ osteocytes did attenuated the cortical bone deficits (Fig. 7B-J). The cortical osteocyte phenotypes, including increased disorganized collagen fibers area, reduced F-actin marked osteocyte networks, altered clustered osteocyte distribution, enlarged osteocyte cell body, impaired SOST^+^ mature osteocytes, and increased cleaved caspase-3^+^ apoptotic cells, were all diminished by injecting MMP14-eGFP virus (Fig. 7B-J). These results suggest that MMP14 is a key downstream protein of ATP6AP2, regulating the cortical osteocyte development.

**Fig 7.**
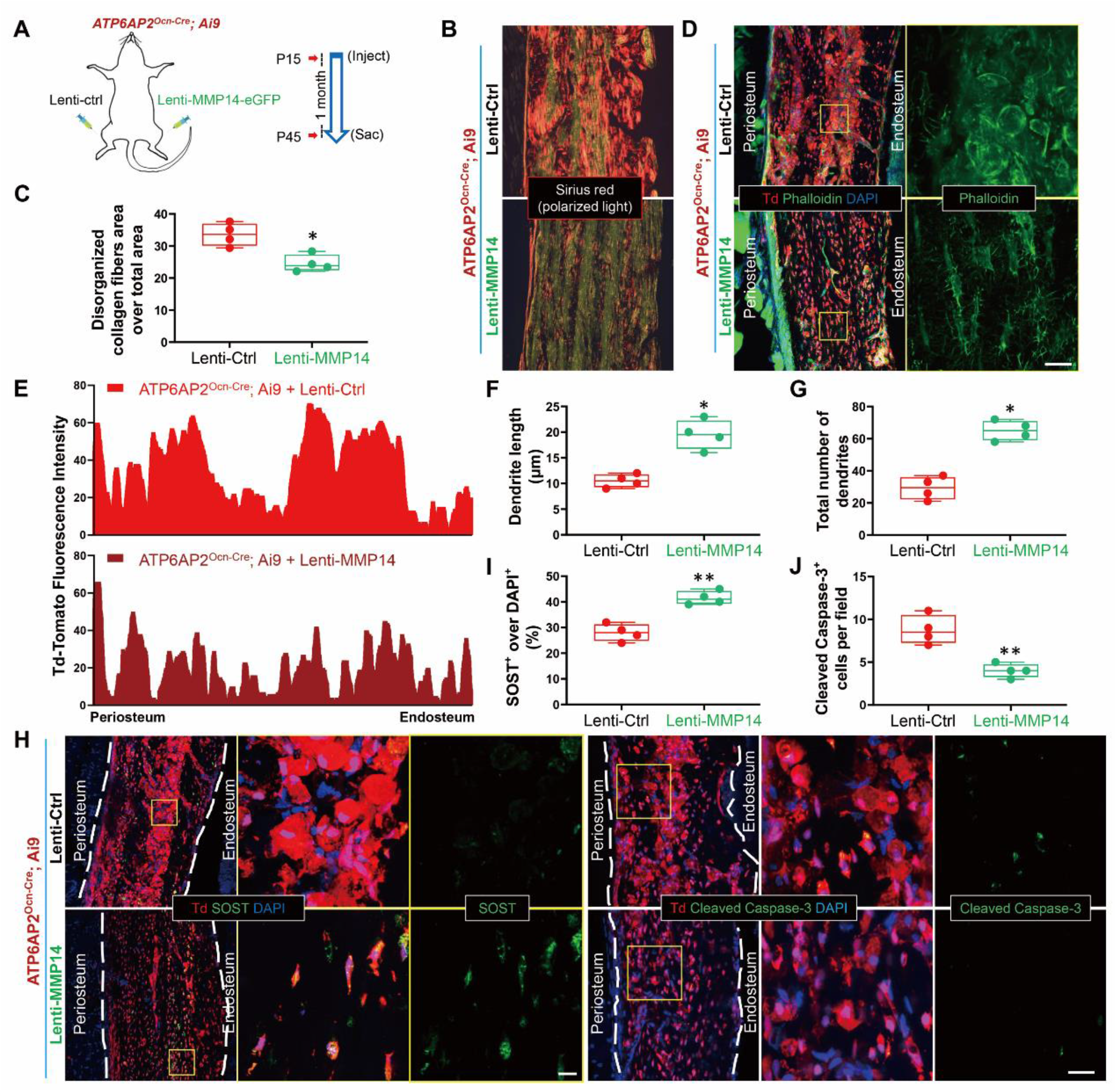
MMP14 attenuation of the cortical osteocyte deficits in ATP6AP2^Ocn-Cre^ mice. **(A)** Ctrl or MMP14-eGFP lentiviral particles were injected into the surface of femur of ATP6AP2^Ocn-Cre^; Ai9 mice at the age of P15. In order to compare the effect of MMP14 overexpression, ctrl or MMP14-eGFP lentiviral particles were injected into different hind legs of the same mouse. Samples were collected 1 month after injection. **(B, C)** Representative images of Sirius red stained cortical bone of indicted mice. Quantification of disorganized collagen distribution rate was shown in C. **(D)** Immunofluorescence staining of dendritic processes and osteocytes with phalloidin-488 for f-actin (green) in femur cortical bone of 1.5-MO ATP6AP2^Ocn-Cre^; Ai9 mice with ctrl or MMP14-eGFP expression. Bar, 20 μm. **(E-G)** Quantification of the TdTomato red fluorescence intensity (E), dendrite length (F) and number of dendrites per osteocyte (G). **(H-J)** Immunofluorescence staining of osteocytes with SOST and cleaved caspase-3 in femur cortical bone of 1.5-MO ATP6AP2^Ocn-Cre^; Ai9 mice with ctrl or MMP14-eGFP expression. Bar, 20 μm. Quantification were shown in I and J. Data are shown as box plots together with individual data points, and whiskers indicate minimum to maximum (n=4 animals). Statistical analysis was performed using unpaired two-tailed t-test. *, P< 0.05. **, P< 0.01

### ATP6AP2 interacting with MMP14 and increasing MMP14 surface targeting

To address how ATP6AP2 regulates MMP14 protein trafficking and surface targeting, we performed co-immunofluorescence staining analyses of MMP14 with endosome/lysosome markers in control and ATP6AP2-KD MC3T3 cells. A significantly reduced MMP14 co-localization with EEA1 (a marker for early endosomes) and Rab7 (a marker for late endosomes), but an increased MMP14 co-localization with LAMP1 (a marker for early lysosomes) were detected in ATP6AP2-KD MC3T3 stable cell line (an OB cell line) (Fig. S7A, B), suggesting a role of ATP6AP2 in suppression of MMP14 targeting to the LAMP1^+^ lysosomes, but not endosomes, and implicating ATP6AP2’s function in preventing MMP14 degradation. We then tested the latter view by examining MMP14’s half-life. Indeed, a faster protein turnover of MMP14 was detected in ATP6AP2-KD MC3T3 cells, exhibiting a half-life of ∼1 hr in ATP6AP2-KD, but ∼2-hr in control cells, (Fig. S7C, D).

To further understand how ATP6AP2 regulates MMP14 surface distribution, we wondered whether both proteins form a complex. To this end, HEK293 cells were co-transfected plasmids encoding MMP14-GFP with plasmids encoding V5 tagged ATP6AP2-full length (FL), and its deletion mutants-the intracellular domain (ICD) (M8.9) and the soluble form or N-terminal domain (ECD) (Fig. 8A). Resulting cell lysates were subjected to immunoprecipitation with anti-MMP14 antibodies. Interestingly, the ATP6AP2-FL and ICD/M8.9, but not the ECD, were detected in the MMP14 immuno-precipitates (Fig. 8B), suggesting that ATP6AP2 interacts with MMP14 via its ICD/M8.9. This view was further supported by the observations of co-localization of MMP14-eGFP with ATP6AP2-FL or ICD/M8.9, but not ECD, by co-immunostaining analysis (Fig. 8C, D). We next examined whether expressing these ATP6AP2-FL and mutant proteins could regulate the cell surface distribution of MMP14-eGFP in MC3T3 cells. As shown in Fig. 8E and F, expressing ATP6AP2-FL and ICD/M8.9, but not the ECD, increased the surface levels of MMP14-eGFP. These results thus suggest a critical role of the ICD domain of ATP6AP2 in forming a complex with MMP14 and in promoting MMP14’s cell surface distribution, supporting the view for ATP6AP2 to promote MMP14 surface targeting by interacting with MMP14 (Fig. 8G).

**Fig 8.**
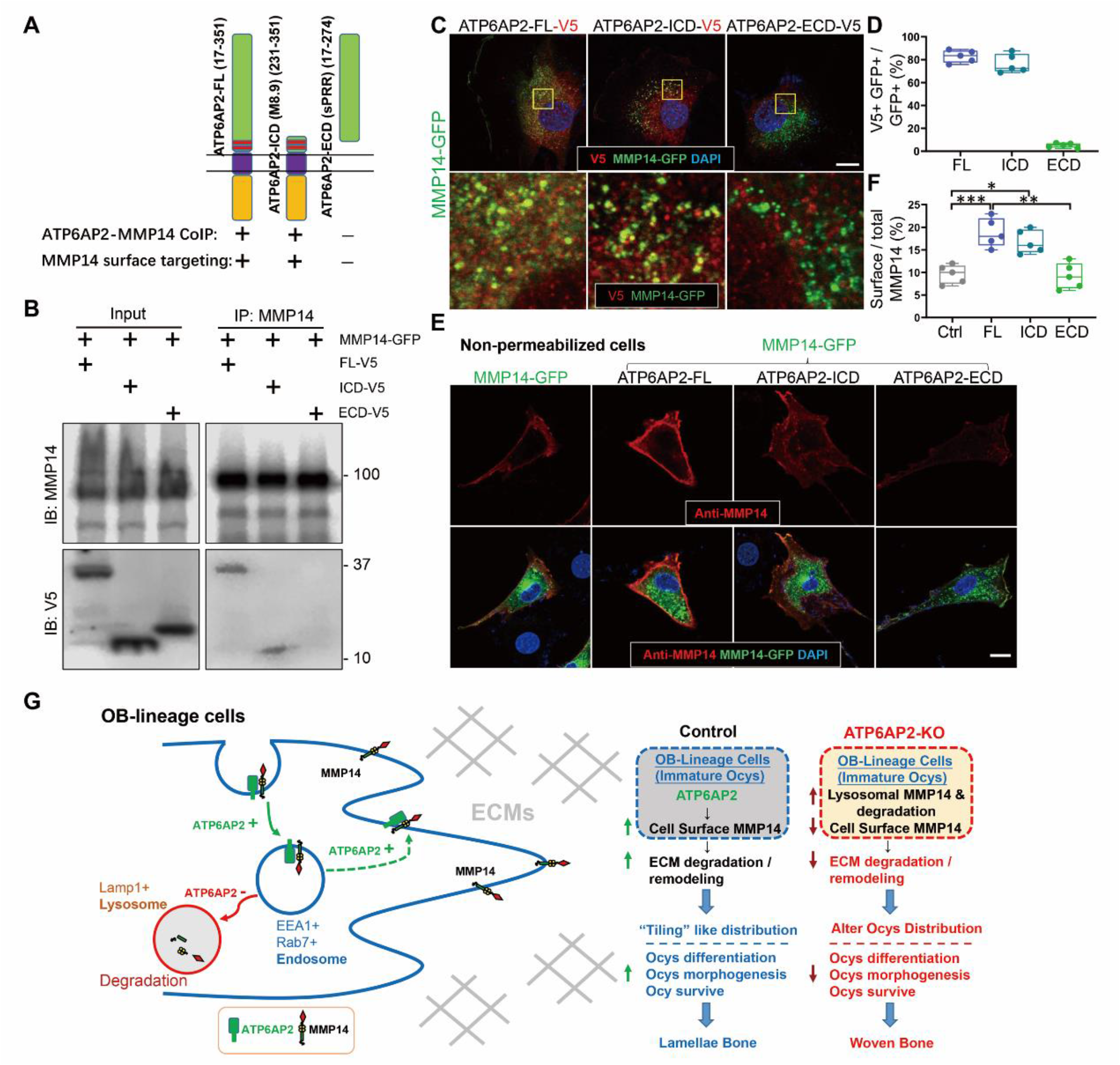
ATP6AP2 increasing MMP14 surface targeting by interacting with MMP14. **(A)** Illustration of various deletion mutants of ATP6AP2. **(B)** Coimmunoprecipitation analysis of ATP6AP2 and its mutant with MMP14. HEK293T cells were transfected with the indicated plasmids. 48 h after transfection, ∼500 µg cell lysates was immunoprecipitated by anti-MMP14 antibodies and A/G agarose. The resulting lysates were subjected to Western blot analysis using indicated antibodies. Approximately 50 µg cell lysates were used as an input. Data presented are representative of three independent experiments. **(C, D)** Co-immunostaining analysis of MMP14-eGFP with various deletion mutants of ATP6AP2 in MC3T3 cells. Representative images are shown in C. Bar, 10 µm. Quantification analysis were shown in D. The co-localization rates of various mutants of ATP6AP2 with MMP14 were determined by the measurement of overlapped signaling (yellow fluorescence) over total GFP^+^ signal. **(E, F)** Immunostaining analysis of cell surface MMP14 in non-permeabilized MC3T3 cells transfected with MMP14-eGFP and various deletion mutants of ATP6AP2. Bar, 10 µm. Quantification analysis of the ratio of cell surface MMP14 over total MMP14 was shown in F. **(G)** Illustration of a working model in which ATP6AP2 deficiency in OB-lineage cells damage MMP14 trafficking in Rab7^+^ vesicle, consequently reducing cell surface levels of MMP14, which leads to disruption of collagen cleavage in osteocytes and ultimately to the loss of formation of osteocyte processes. Data in (D) and (F) are shown as box plots together with individual data points, and whiskers indicate minimum to maximum (5-different assays). P values obtained by one-way ANOVA followed by Tukey post hoc test. *, P< 0.05. **, P< 0.01. ***, P< 0.001.

## Discussion

The present study provides evidence for ATP6AP2 in Ocn-Cre^+^ OB-lineage cells to be critical for OB-to-osteocyte transition and thus osteocyte development, distribution and network building. This study also uncovers ATP6AP2’s functions in promoting lamellae bone formation likely by increasing MMP14 surface targeting and activation in immature osteocytes. Further mechanistic studies lead to a working model that is depicted in Fig. 8G. In this model, ATP6AP2 increases MMP14 surface targeting largely from endosomes by interacting with MMP14, and thus enhances MMP14 mediated ECM degradation and remodeling, osteocyte precursor cells’ distribution, osteocytes differentiation, morphogenesis and survive. This study not only identifies ATP6AP2 as a key regulator of MMP14, but also reveal a pathway from ATP6AP2 to MMP14 in promoting osteocyte development.

ATP6AP2 is known to play important roles in multiple signaling pathways and in multiple organs (Ichihara & Yatabe, 2019). In addition to function as a PRR, inducing the conversion of angiotensinogen to angiotensin I and regulating blood pressure (Nguyen et al., 2002), it is a critical regulator of Wnt/β-catenin signaling and V-ATPase by its interaction with Wnt receptors (e.g., LRP6 and frizzled) and V-ATPase membrane proteins (Buechling et al., 2010; Cruciat et al., 2010). By its interaction with LRP6 and frizzled, ATP6AP2 promotes Wnt/β-catenin signaling (Buechling et al., 2010; Cruciat et al., 2010). In line with this view is our proteomic analysis, which identified reduced LRP6 surface targeting in ATP6AP2 KO OB-lineage cells (Fig. 5B, C). However, in osteocytes of the cortical bone regions, we speculate that ATP6AP2-induced MMP14 surface targeting and activation may be a key pathway to regulate cortical bone ECM remodeling and osteocyte development.

How does ATP6AP2 promotes MMP14 surface targeting? Encouraged by the following observations, we propose that ATP6AP2 promotes MMP14 surface targeting by interacting with MMP14 largely in endosomes. First, ATP6AP2 forms a complex with MMP14 via its C-terminal ICD (Fig. 8A-D). This ICD domain, also called M8.9, binds to the V-ATPase complex proteins and promotes V-ATPase complex assemble and activation (Ludwig et al., 1998). Second, in ATP6AP2-KD MC3T3 cells, expression of this ICD domain is sufficient to increase MMP14 surface targeting (Fig. 8E, F). Third, in addition to the reduced MMP14 surface targeting in ATP6AP2-KD MC3T3 cells, MMP14 in EEA1^+^ or Rab7^+^ endosomes were decreased, but MMP14 in Lamp1^+^ lysosomes were increased, in the mutant cells (Fig. S7A, B). Finally, similar V-ATPase’s complex and cell type specific effects on MMPs’ activities are also observed in different tumor cells (Chung et al., 2011; Hoshino et al., 2012; Smith et al., 2016). Given our results that the surface and total levels of MMP14 were decreased in ATP6AP2 KO cells (Fig. 5), we speculate that ATP6AP2 may promote V-ATPase driven vesicle pH and MMP14 surface targeting, but not V-ATPase dependent MMP14 degradation. In line with the view for vesicle pH to be critical for MMP14 surface targeting is the report that CLIC4, a chloride intracellular channel 4 that regulates vesicular HCl and pH, colocalizes with MMP14 in the late endosomes, and promotes MMP14 surface targeting and activation in retinal pigment epithelium (RPE) (Hsu et al., 2019). In addition, ATP6AP2 deficiency was also reported to impair V-ATPase assembly and subsequent defects in glycosylation and autophagy (Rujano et al., 2017). Further investigation is needed to address whether ATP6AP2 affects the glycosylation of MMP14.

As a subunits of V-ATPase, ATP6AP2 has been reported to contribute to various pathological events/diseases, such as fibrosis, hypertension, acute kidney injury, cardiovascular disease, cancer, obesity, and various other diseases (Ichihara & Yatabe, 2019). Our study demonstrates a critical role of ATP6AP2 in promoting osteocyte development. We speculate that ATP6AP2, as a key regulator of MMP14, prevents the woven bone accumulation likely by promoting bone ECM remodeling and osteocyte development. Dysregulation of ATP6AP2 to MMP14 pathway may underlie the pathogenesis of many ATP6AP2/MMP14 involved disorders. We hope to further test this view in future.

## Materials and Methods

### Animals and Reagents

The ATP6AP2^flox/X^ mice (kindly provided by Dr. Frederique Yiannikouris, University of Kentucky; and Dr. Genevieve Nguyen, INSERM, France) were crossed with osteocalcin (Ocn)-Cre (kindly provided by Dr. Tom Clemens, Johns Hopkins Medical School) or Dmp1-Cre (Jackson lab, Stock No 023047) transgenic mice to generate OB- or Osteocyte-selective conditional knockout (CKO) mutant mice, ATP6AP2^Ocn-Cre^ or ATP6AP2^DMP1-Cre^, respectively. The mutant mice were in C57BL/6J mouse background (for more than 6 generations). The Ai9 mice were purchased from the Jackson Laboratory (Stock No 007909), which are in C57BL/6J background. All experimental procedures were approved by the Institutional Animal Care and Use Committee at Case Western Reserve University (IACUC, 2017-0121 and 2017-0115), according to US National Institutes of Health guidelines.

Mouse monoclonal antibodies, including β-actin (A1978, Sigma), GAPDH (NB300-221, Novus), Rab7 (ab50533, Abcam), SOST (AF1589, R&D) and V5 (V8012, Sigma-Aldrich), were used. Rabbit monoclonal antibodies Ki67 (ab16667, Abcam), MMP14 (ab51074, Abcam), Runx2 (ab192256, Abcam) and Lrp1 (ab92544, Abcam) were used. Rabbit polyclonal antibody Cleaved Caspase-3 (9661, Cell signaling technology), ATP6AP2 (HPA003156, Sigma Aldrich) and EEA1 (ab2900, Abcam) were used. Rat Lamp1 (1D4B-c, DSHB) antibody and Syrian hamster monoclonal E11(gp38 or gp36, ab11936, Abcam) antibody were used. Alexa Fluor 488 phalloidin (f-actin, A12379) was purchased from Thermo Fisher. Secondary antibodies were purchased from Jackson ImmunoResearch Laboratories, Inc. Calcein (C0875) and cycloheximide solution (CHX, C4859) were obtained from Sigma-Aldrich. Other chemicals and reagents used in this study were of analytical grade.

### Plasmids and lentiviruses

ATP6AP2-V5-FL and ATP6AP2-V5-ICD plasmids were purchased from DNASU (ATP6AP2 in pLX304, HsCD00446844 and HsCD00437681). ATP6AP2-V5-ECD was made by cloning the ECD domain (17-274) of ATP6AP2 into the pLX304 plasmid (KpnI and XbaI). MMP14-eGFP plasmid was purchased from Addgene (item ID 89819). Renin Receptor shRNA Lentiviral Particles (shR-PRR) was obtained from Santa Cruz (sc-62935-V). The authenticity of all constructs was verified by DNA sequencing.

The 3rd generation cre-dependent lentiviral system was used to generate the pLV-FLEX-MMP14-eGFP lentiviral particle. MMP14-eGFP was cloned into pLV-FLEX lentiviral plasmid. The empty ctrl or MMP14-eGFP lentiviral plasmid (7.5 μg) was co-transfected into 293 cells with packaging plasmid pRSV-Rev (5 μg, Addgene #12253), pMDLg/pRRE (2.5 μg, Addgene #12251), and envelope plasmid pMD2.G (0.8μg, Addgene #12259) to package the lentivirus. The lentivirus was concentrated before use.

### In vitro OB/OC-lineage cell cultures

Whole bone marrow cells were flushed from long bones of Ctrl and ATP6AP2-deficient mice and plated on 100 mm culture plates in DMEM containing 1% penicillin/streptomycin (P/S), 10% fetal bovine serum (FBS) for 2 days. For OB-lineage culture, plates with adherent cells were replaced with fresh culture medium every 3 days. After 7 days, passaging cells (BMSCs) by trypsin digestion, 1×10^4^/cm^2^ were plated for experiments.

For OC-lineage culture, non-adherent cells were harvested and subjected to Ficoll-Hypaque gradient centrifugation for purification of BMMs. Cells were plated on 100 mm culture dishes in α-MEM containing 10% FBS, 1% P/S and 10 ng/ml recombinant M-CSF. For osteoclastogenesis, 5×10^4^ BMMs were incubated with OC differentiation medium containing 10 ng/ml recombinant M-CSF and 100 ng/ml recombinant RANKL. Mature OC began to form on day 4 to 5 after RANKL treatment. The cells were then subjected to TRAP staining to confirm their OC identity.

Primary OB cultures were prepared from long bones of mice. Briefly, small bone pieces were incubated in collagenase solution to remove all remaining soft tissue and adhering cells, then transfer to 60 mm culture dishes containing DMEM medium supplemented with 10% FBS, 1% penicillin/streptomycin, 10 mM β-glycerophosphate and 50 μM L-ascorbic acid-2-phosphate. Replace culture medium three times per week. Bone cells will start to migrate from the bone chips after 3-5 days. After two weeks, the monolayer is trypsinized by incubating the cells with trypsin solution. OBs were plated on 100 mm tissue culture plates in α-MEM containing 10% fetal bovine serum (FBS), 1% penicillin/streptomycin (P/S).

Primary osteocyte cultures were isolated from mouse long bones as described previously (Stern et al., 2012). In brief, long bones (femur and tibia) were aseptically dissected from skeletally mature 3-MO mice. Collagenase solution was prepared as 300 active U/mL collagenase type-IA (Sigma-Aldrich, St. Louis, MO, USA) dissolved in DMEM. EDTA tetrasodium salt dehydrate (EDTA) solution (5 mM, pH = 7.4; Sigma-Aldrich) was prepared in magnesium and calcium-free Dulbecco’s phosphate-buffered solution (DPBS; Mediatech) with 1% BSA (Sigma-Aldrich). All steps of the digestion took place in 5 mL solution in a 6-well plate, on a rotating shaker set to 200 RPM, in a 37°C and 5% CO_2_ humidified incubator. Following each sequential digestion, the digest solution with suspended cells was removed from the bone pieces and kept. The combined cell suspension solution was spun down at 900 rpm for 5 min, the supernatant was removed from the cell pellet, and cells were resuspended in culture medium and counted.

### Cell line and transfection

MC3T3-E1 or HEK293 cells were maintained in DMEM supplemented with 10% fetal calf serum and 1% penicillin/streptomycin. For transient transfection, MC3T3-E1 cells were plated at a density of 10^6^ cells per 10-cm culture dish and allowed to grow for 12 hours before transfection by using Lipofectamine™ 3000 Transfection Kit (L3000, Invitrogen). 48 hours after transfection, cells were subjected to immunostaining analysis. HEK293 cells were transfected by Polyethylenimine (PEI), as described previously (L. Xiong et al., 2015; Xiong et al., 2016). In brief, 12 μg DNA mixture was prepared in serum-free DMEM, 6 μl PEI (based on a 3:1 ratio of PEI: total DNA) was added to the diluted DNA and mixed immediately by pipetting. After incubation for 20 min at room temperature, the DNA/PEI mixture was added to cells, and 48 hours later, transfected cells were subjected to Western blot or CO-IP assay.

The ATP6AP2-KD cell line was obtained by infection of MC3T3-E1 cells with lentiviral particles encoding scramble control or shRNA-ATP6AP2, respectively. In brief, cells were infected with the lentiviral particles for 1 day in polybrene (2 μg/ml) medium. At day 3, the culture medium was removed and replaced with complete medium (without polybrene). After 5-6 days, stable clones expressing the shRNA were selected via puromycin dihydrochloride (5 μg/ml) that induces death of the un-transduced cells.

### Cell lysis, western blot, and co-immunoprecipitation

Cells were lysed in lysis buffer containing 50 mM Tris-HCl (pH7.5), 150 mM NaCl, 1%(v/v) Triton X-100, 0.1% SDS, 0.5% deoxycholate and 1 mM EDTA, supplemented with protease inhibitors (1 μg/mL leupeptin and pepstatin, 2μg/mL aprotinin and 1 mM PMSF) and phosphatase inhibitors (10 mM NaF and 1 mM Na3VO4). Whole cell extracts were fractionated by SDS-PAGE and transferred to a nitrocellulose membrane (Bio-Rad). After incubation with 5% milk in TBST (10 mM Tris, 150 mM NaCl, 0.5% Tween 20, pH 8.0) for 1-h, the membrane was incubated with indicated antibodies overnight at 4°C. Membranes were washed with TBST for three times and incubated with a 1:5000 dilution of horseradish peroxidase-conjugated anti-mouse or anti-rabbit antibodies for 1-h. Blots were washed with TBST three times and developed with the ECL system (Bio-Rad). Immunoprecipitation was performed with an anti-MMP14 antibody. Western blotting was performed using the ECL procedure according to the manufacturer’s instructions (Bio-Rad), with a mouse monoclonal V5 antibody to detect the co-immunoprecipitation ATP6AP2-V5 or anti-MMP14 rabbit polyclonal antibody for MMP14 proteins.

### Immunofluorescence staining and imaging analysis

For bone section staining, mouse femur, vertebrate bones or skull bones were fixed with 4% PFA in PBS at 4°C overnight, then decalcified in 14% EDTA for 10 days. The samples were place in OCT and cut into 50 μm sections on a cryostat (HM550, Thermo Scientific). For cell immunofluorescence staining, cells on coverslips were fixed with 4% paraformaldehyde at room temperature for 20 min. Bone sections or cells were permeabilized with 0.2% Triton X-100 for 8 min, and then subjected to co-immunostaining analysis using indicated antibodies. For cell surface protein staining, cells were directly subjected to co-immunostaining analysis without Triton X-100 (non-permeabilized). Stained sections or cells were washed 3 times with PBS and mounted with VECTASHIELD (H-1500; Vector Laboratories) and imaged by Confocal Microscope at room temperature. Fluorescent quantification was performed using ZEN software according to the manufacturer’s instructions (Carl Zeiss).

### Micro-computed tomography (μCT)

The μCT analyses were carried out as described previously (Xiong et al., 2017). Excised femurs from mice were scanned using the Scanco µCT40 desktop cone-beam micro-CT scanner (Scanco Medical AG, Brüttisellen, Switzerland using µCT Tomography v5.44). Scans were automatically reconstructed into 2-D slices and all slices were analyzed using the µCT Evaluation Program (v.6.5-2, Scanco Medical). The femur was placed inverted in a 12mm diameter scanning holder and scanned at the following settings: 12µm resolution, 55kVp, 145µA with an integration time of 200ms. For the cortical analysis, the bone was scanned at the midshaft of the bone for a scan of 25 slices. The region of interest (ROI) was drawn on every slice and fitted to the outside of the cortical bone, to include all the bone and marrow. The threshold for cortical bone was set at 329. The 3-D reconstruction (µCT Ray v3.8) was performed using all the outlined slices. Data were obtained on bone volume (BV), total volume (TV), BV/TV, bone density and cortical thickness.

### Bone histomorphometric analysis

Bone histomorphometric analyses were carried out as previously described (Schmitz, Laverty, Kraus, & Aigner, 2010; Xiong et al., 2020). In brief, mouse tibia and femurs were fixed overnight in 10% formalin, decalcified in 14% EDTA, embedded in paraffin, sectioned, and subjected for hematoxylin and eosin (H&E), Safranin-O analyses which were counterstained by fast green and Picro-Sirius red staining. Brightfield and polarized microscopic images were obtained (BZ-X710 microscope with a pair of circular polarizing filters, Keyence) and analyzed using ImageJ. For the picro-sirius red staining, polarized images were used to measure woven bone area and brightfield images were used to measure cortical area. In brightfield microscopy collagen is red on a pale yellow background. Under polarized light, the larger collagen fibers are deep red, bright yellow or orange, and the thinner ones are green.

### Dynamic bone histomorphometry to measure the rate of bone formation in vivo

Briefly, mice (P30) were injected (intraperitoneally) with fluorochrome-labeled calcein green (10 mg/kg, Sigma–Aldrich) twice (5 days interval). The mice were sacrificed 2 d after the second injection. The femurs were fixed in 70% (vol/vol) ethanol overnight, embedded in methyl methacrylate, and sectioned at 7–10 μm. Images were obtained using Zeiss LSM 800 fluorescence microscope. The mineral apposition rate (MAR) in μm/d and bone formation rate (BFR) [BFR = MAR × MS (mineral surface) / BS (bone surface)] were calculated from fluorochrome double labels at the periosteum and endosteum surfaces.

### Cell surface biotinylation and Proteomic analysis

Cultured OBs were incubated with Sulfo-NHS-biotin (ThermoFisher Scientific) in PBS for 45 min at 4 °C, and treated with 10 mM glycine for 20 min at 4 °C to terminate the crosslinking reaction. After rinse with PBS, cells were lysed in RIPA buffer. Biotinylated proteins were precipitated overnight with avidin agarose beads (Pierce). Finally, beads were washed 3 times with PBS, and 1X SDS loading buffer was added to collect biotinylated surface proteins.

For proteomic assay, biotinylated surface proteins were separated by SDS-Page gel and submitted for LC-MS/MS analysis. The LC-MS system was a ThermoScientific Orbitrap Elite mass spectrometer system. The HPLC column was a Dionex 15 cm x 75 μm id Acclaim Pepmap C18, 2μm, 100 Å reversed-phase capillary chromatography column. 5 μL volumes of the extract were injected and the peptides eluted from the column by an acetonitrile/0.1% formic acid gradient at a flow rate of 0.3 μL/min were introduced into the source of the mass spectrometer on-line. The microelectrospray ion source is operated at 2.5 kV. The digest was analyzed using the data dependent multitask capability of the instrument acquiring full scan mass spectra to determine peptide molecular weights and product ion spectra to determine amino acid sequence in successive instrument scans. The data were analyzed by using all CID/HCD spectra collected in the experiment to search the mouse UniProtKB database with the search Sequest. The protein and peptide identifications were validated with the program scaffold.

### Collagen-degradative activity

The culture dishes were coated with type I collagen according to the instructions (A10483-01, Gibco). In brief, the rat tail Collagen I (3 mg/mL) were mixed with dH_2_O, 1N NaOH, and 10 X PBS to achieve at pH of 6.5-7.5. The mixed Collagen I were then dispensed into the desired dishes immediately and incubate at 37°C in humidified incubator for 30–40 minutes or until a firm gel is formed. The dishes was rinsed with sterile 1 X PBS and labeled with the fluorescent dye (Alexa fluor 488 or 594 NHS Ester) as described previously (Doyle, 2018). Primary cultured OBs or osteocytes were added into the dishes and protected from light for 1 to 3 days. Images were obtained using Zeiss LSM 800 fluorescence microscope.

### Lentiviral particle injection *in vivo*

The p15 mice were placed under general anesthesia by i.p.injection of ketamine and xylazine. The femur areas were shaved and cleaned with iodine and ethanol solutions for aseptic skin preparation. A fine glass pipette (diameter of tip being 20-40 μm) was inserted into the area between muscle and femur to deliver ctrl or the pLV-FLEX-MMP14-eGFP lentiviral particle to the surface of femur. After delivery, the glass pipette was pulled out slowly 4-5 min after injection. 1 month after lentivirus delivery, the mice were euthanized and tissue were collected.

### RNA isolation and real time-PCR

Total RNA was isolated by Trizol extraction (Invitrogen, Carlsbad, CA, USA). Q-PCR was performed by using Quantitect SYBR Green PCR Kit (Bio-Rad) with a Real-Time PCR System (Opticon Monitor 3). MMP14 primers (5′-AGTGCCCTATGCCTACATCC-3′ and 5′-GGAACCCTCCTTCACCATCA-3′) and β-actin primers (5′-AGGTCATCACTATTGGCAACGA-3′ and 5′-CATGGATGCCACAGGATTCC-3′) were used.

### Statistical analysis

For in vivo studies, 3-6 mice per genotype per assay were used. For in vitro cell biological and biochemical studies, each experiment was repeated at least 3 times. Statistical analyses were performed with GraphPad Prism 8 using unpaired student’s two-tailed t-test, one-way ANOVA with Tukey post hoc test, or two-way ANOVA with Bonferroni post doc test. In all experiments, the significance level was set at P< 0.05 (* P< 0.05, ** P< 0.01, *** P< 0.001).

## Acknowledgements

We thank Drs. Genevieve Nguyen (INSERM, France) and Frederique Yiannikouris (University of Kentucky) for providing us the ATP6AP2^flox/flox^ mice, Ms. Xue-Mei Cao and Dr. Maria S Johnson (University of Alabama at Birmingham) for μCT analysis, and members of the Xiong and Mei laboratories for helpful discussions. This study was supported in part by grants from the National Institutes of Health (AG045781, AG051510, and AG066526) (to WCX).

## Author Contributions

LX and WCX designed and performed experiments and data analyses. HG, JP, and DL performed some experiments. XR assisted with data analyses. LM and WCX supervised the experiments and assisted with data analyses. LX and WCX wrote the manuscript.

## Supplementary figures

**Fig S1.**
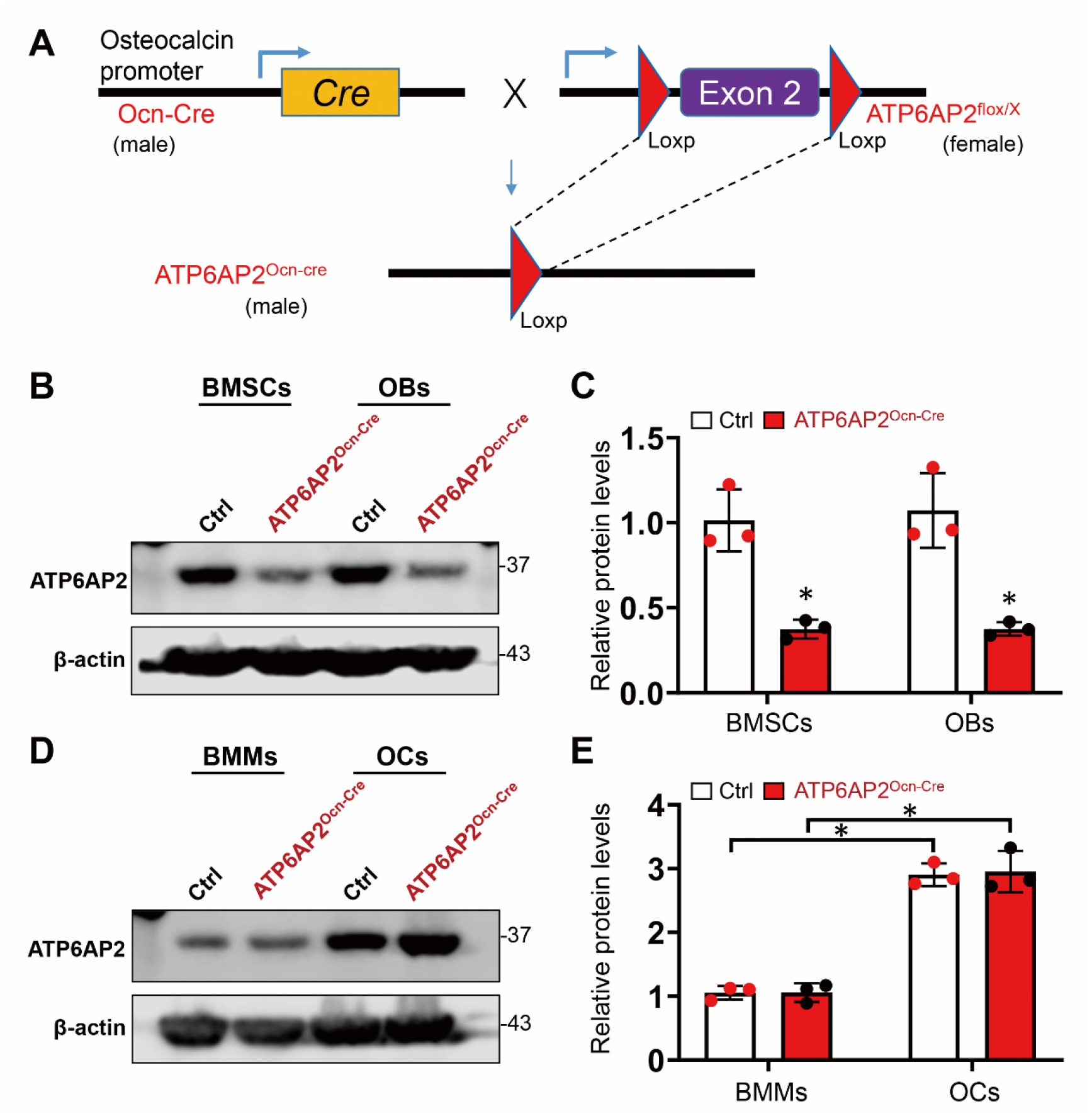
Generation of OB-selective ATP6AP2 KO mice. **(A)** Strategy to cleave exon 2 of *ATP6AP2* flanked by loxP sites in the ATP6AP2^flox^ allele. As *ATP6AP2* is on the X chromosome, ATP6AP2^flox/X^ female mice were crossed with male osteocalcin (Ocn)-Cre transgenic mice to generated OB-selective conditional knockout mutant mice, ATP6AP2^Ocn-Cre^. **(B)** Western blotting analysis of ATP6AP2 expression in primary cultured BMSCs and OBs from 3-MO Ctrl and ATP6AP2^Ocn-Cre^ mice. β-actin was used as the loading controls. **(C)** Quantification analysis of B. **(D)** Western blotting analysis of ATP6AP2 expression in primary cultured BMMs and OCs from 3-MO Ctrl and ATP6AP2^Ocn-Cre^ mice. **(E)** Quantification analysis of D. Data in (C) and (E) are presented as mean± SD (n=3 independent experiments). *, P< 0.05 as determined by two-way ANOVA with Bonferroni post hoc analysis for multiple comparisons test.

**Fig S2.**
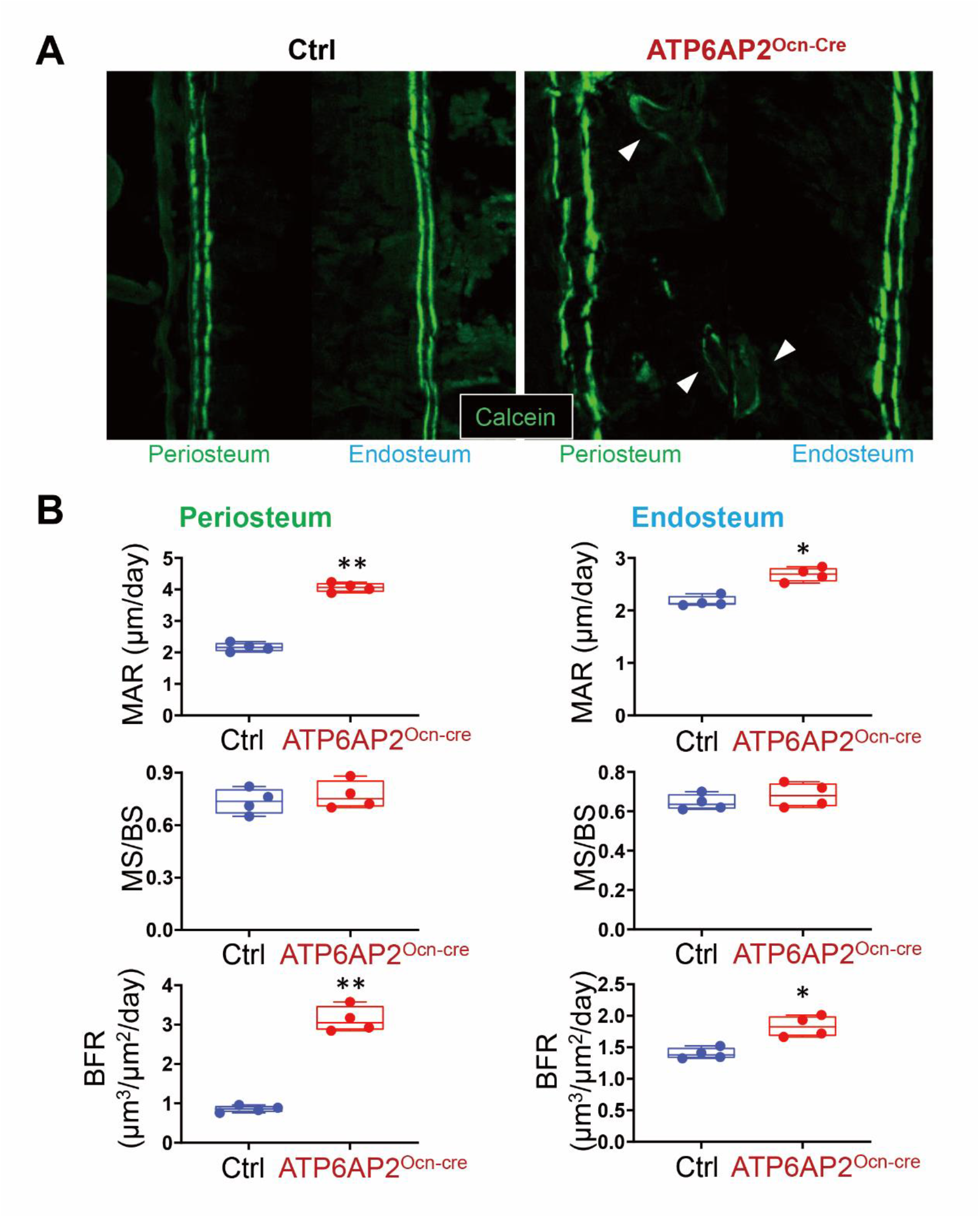
Increased bone formation rate in the cortical bone of ATP6AP2^Ocn-Cre^ mice. **(A)** Representative images of histologic sections showing calcein labeling of periosteum and endosteum bone regions in femur of ctrl and ATP6AP2^Ocn-Cre^ mice at age of 1-MO. Note that there are calcein labeled mineralized areas in the center of the cortical bone of ATP6AP2 ^Ocn-Cre^ mice (white arrowhead). **(B)** MAR (mineral apposition rate), MS (mineral surface) / BS (bone surface), and BFR (bone formation rate) are presented. Data in (B), (F) and (H) are shown as box plots together with individual data points, and whiskers indicate minimum to maximum (n=4 animals per genotype). P values obtained by unpaired two-tailed t-test. *, P< 0.05. **, P< 0.01.

**Fig S3.**
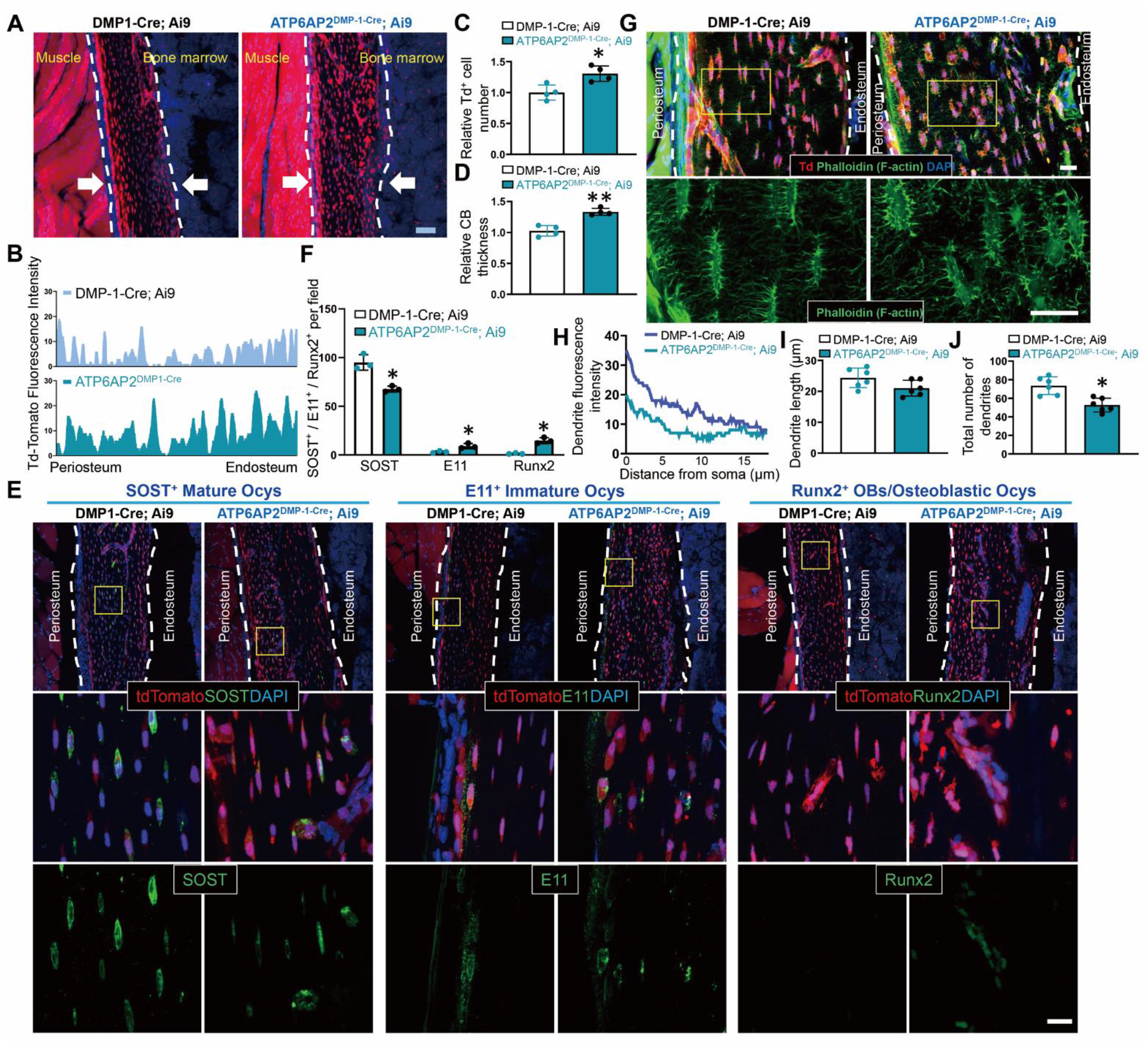
Similar, but weaker, bone deficit in ATP6AP2 ^DMP-1-Cre^ mice. **(A)** Td-Tomato expression in the cortical bone region of femur sections from 1-MO mice with targeted deletion of ATP6AP2 in cells expressing Dmp-1 (TdTomato^+^, ATP6AP2^Dmp-1-Cre^; Ai9) and their littermates (Dmp-1-Cre; Ai9). Bar, 50 μm. **(B-D)** Quantifications of the TdTomato red fluorescence intensity (B), the tdTomato^+^ cell density (C) and cortical bone thickness (D). **(E)** Immunofluorescence of femur cortical bone using SOST (mature osteocytes marker), E11 (early, immature osteocyte marker) and Runx2 (preosteoblast, osteoblast, osteoblastic osteocyte marker) antibodies. Bar, 20 μm. **(F)** Quantification of the percentage of SOST^+^, E11^+^ and Runx2^+^ cells per field. **(G)** Immunofluorescence staining of osteocytes and dendritic processes with phalloidin-488 for f-actin (green) in femur cortical bone of 1-MO DMP-1-Cre; Ai9 and ATP6AP2^DMP-1-Cre^; Ai9 mice. Bar, 20 μm **(H-J)** Quantification of dendrite fluorescence intensity (H), dendrite length (I) and number of dendrites per osteocyte (J). Data in (C), (D), (F), (I) and (J) are presented as mean± SD (n = 3, 4 or 6 animals per genotype). *, P< 0.05, **, P< 0.01 as determined by unpaired two-tailed t-test.

**Fig S4.**
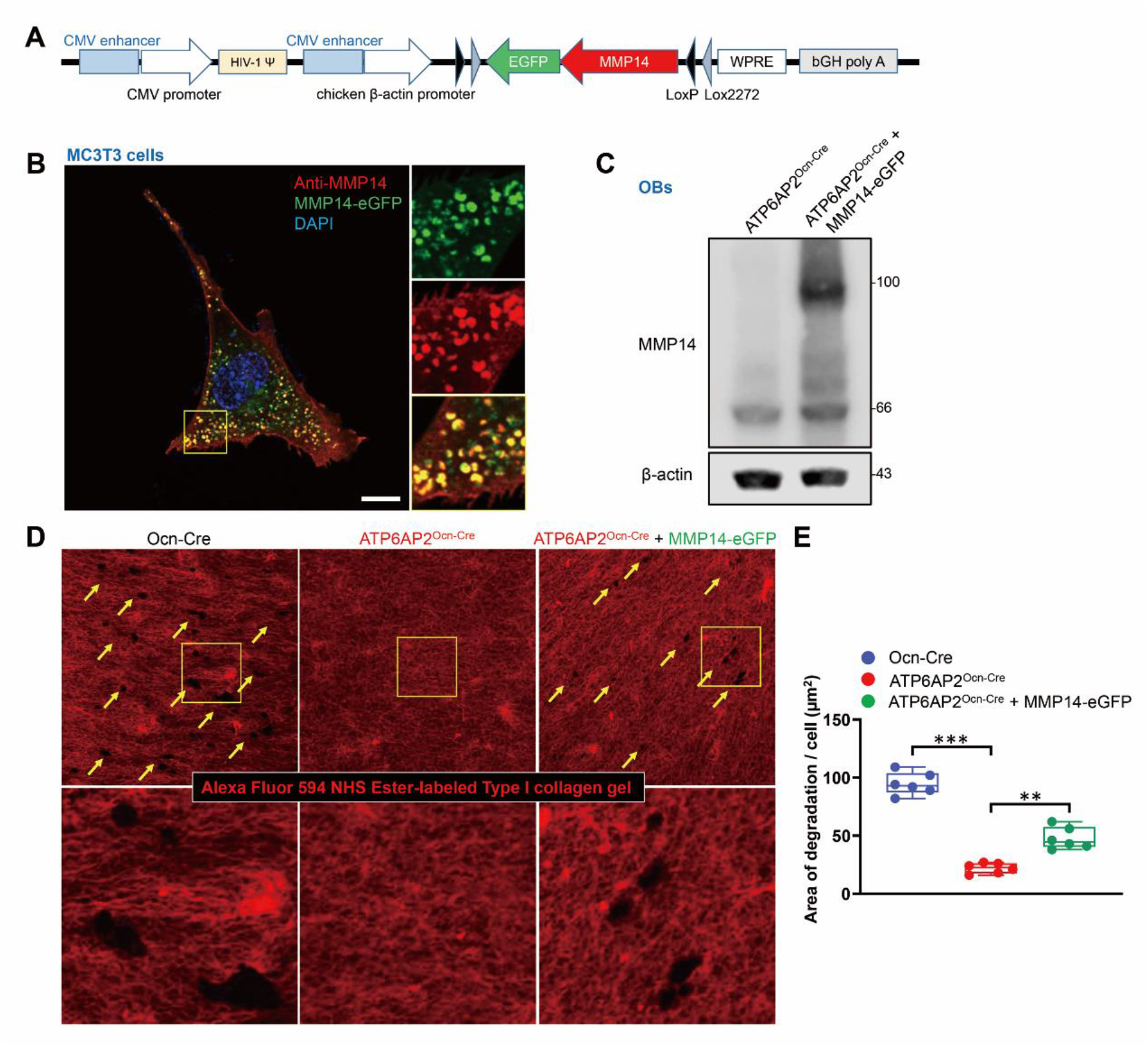
Overexpression of MMP14 promotes collagen-degradative activity of ATP6AP2-KO OBs. **(A)** Generation of MMP14-eGFP lentiviral particles. **(B)** Immunofluorescence analysis of MMP14-eGFP lentivirus plasmid expression in MC3T3 cells. Cells were co-transfected with MMP14-eGFP lentivirus and cre plasmids. Bar, 10 μm. **(C)** Western blot analysis of MMP14-eGFP, which was increased in ATP6AP2^Ocn-Cre^ + MMP14-eGFP OBs. OBs were infected with lentivirus and then purified by FACS. **(D)** Confocal micrographs showing degradation of the fluorescent type I collagen layer, indicated by dark holes, by ctrl, ATP6AP2-KO and ATP6AP2-KO with MMP14-eGFP overexpression OBs. The type I collagen gel was labeled by Alexa fluor 594 NHS ester dye before seeding the cells. **(E)** Quantitative measurement in D. Data are shown as box plots together with individual data points, and whiskers indicate minimum to maximum (n=6 independent experiments). P values obtained by one-way ANOVA followed by Tukey post hoc test. **, P< 0.01. ***, P< 0.001.

**Fig S5.**
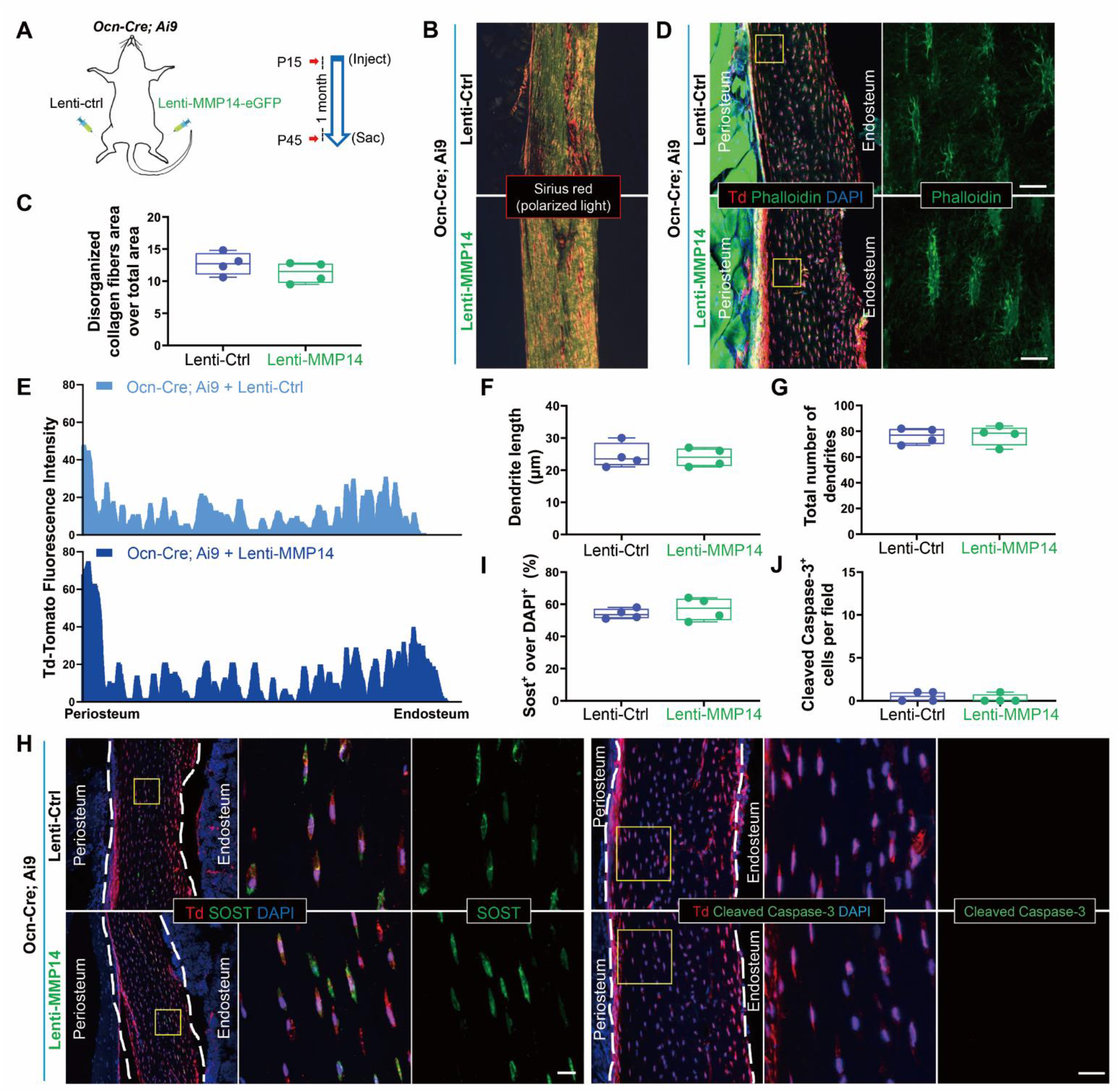
No significant changes in femur of ctrl mice expressing MMP14-eGFP fusion protein. **(A)** Ctrl or MMP14-eGFP lentiviral particles were injected into the surface of femur of Ocn-Cre; Ai9 mice at the age of P15. In order to compare the effect of MMP14 overexpression, ctrl or MMP14-eGFP lentiviral particles were injected into different hind legs of the same mouse. Samples were collected 1 month after injection. **(B, C)** Representative images of Sirius red stained cortical bone of indicted mice. Quantification of disorganized collagen distribution rate was shown in C. **(D)** Immunofluorescence staining of dendritic processes and osteocytes with phalloidin-488 for f-actin (green) in femur cortical bone of 1.5-MO Ocn-Cre; Ai9 mice with ctrl or MMP14-eGFP expression. Bar, 20 μm. **(E-G)** Quantification of the TdTomato red fluorescence intensity (E), dendrite length (F) and number of dendrites per osteocyte (G). **(H-J)** Immunofluorescence staining of osteocytes with SOST and cleaved caspase-3 in femur cortical bone of 1.5-MO Ocn-Cre; Ai9 mice with ctrl or MMP14-eGFP expression. Bar, 20 μm. Quantification were shown in I and J. Data are shown as box plots together with individual data points, and whiskers indicate minimum to maximum (n=4 animals). Statistical analysis was performed using unpaired two-tailed t-test.

**Fig S6.**
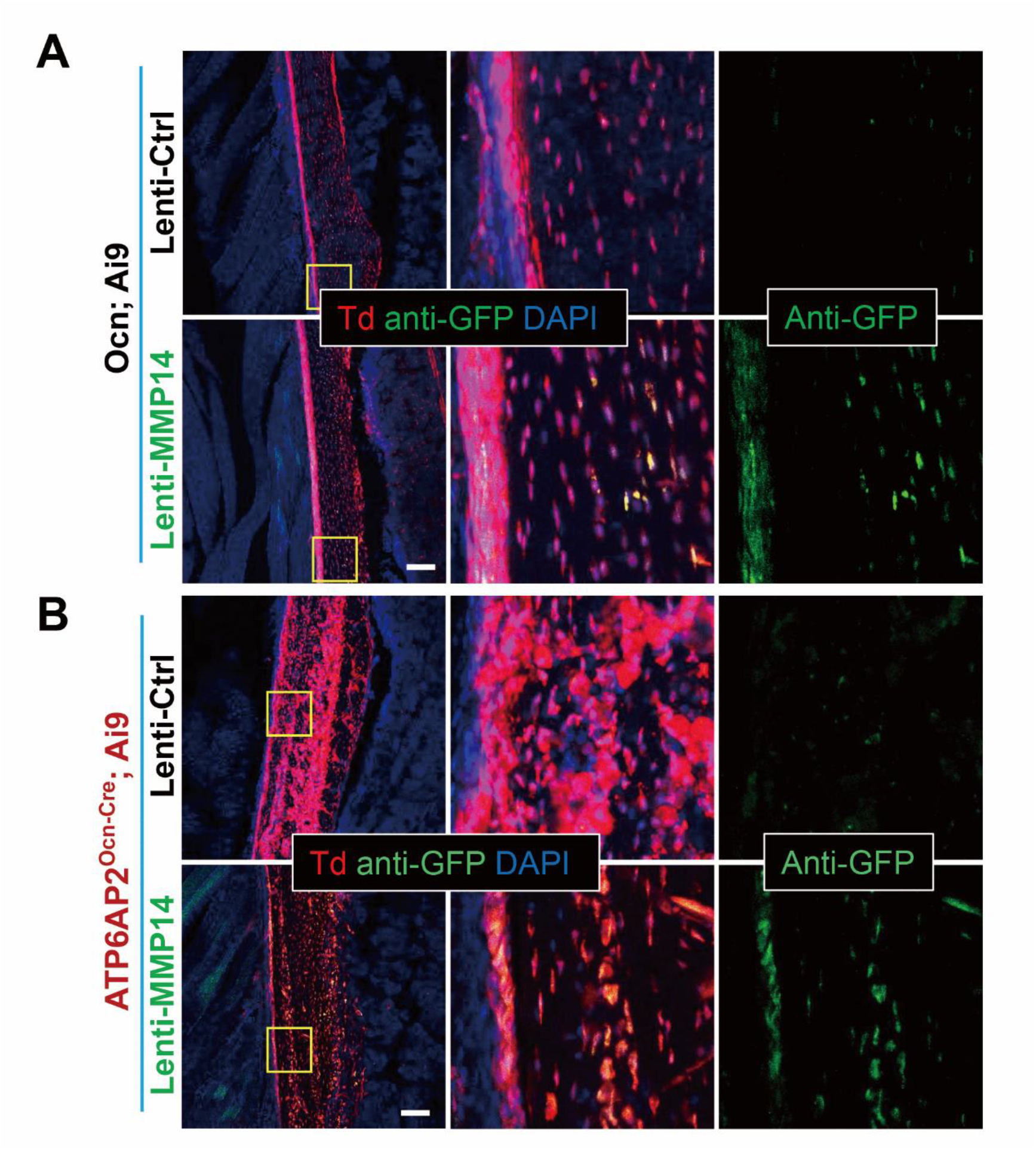
Expression of MMP14-eGFP in femur of mice. **(A, B)** Immunofluorescence analysis of MMP14-eGFP expression in femur of indicted mice. Bar, 100 μm.

**Fig S7.**
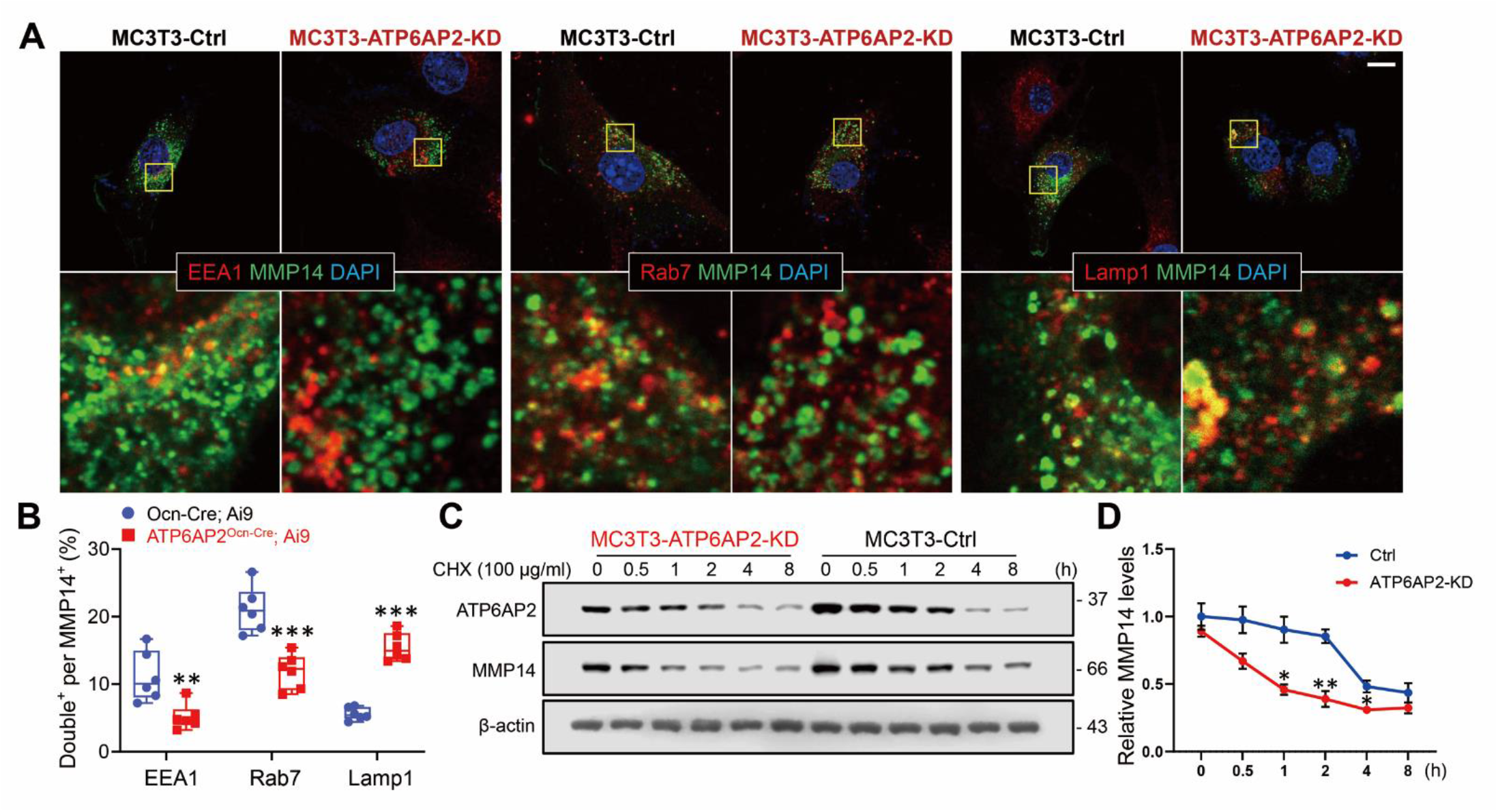
ATP6AP2 regulates MMP14 protein stability. **(A-B)** Co-immunostaining analysis of MMP14 with EEA1 (a marker for early endosomes), Rab7 (a marker for late endosomes) or Lamp1 (a marker for late endosomes and early lysosomes) in control and ATP6AP2-KD MC3T3 cells. Representative images were shown in A. Images marked with yellow squares were amplified and shown in the bottom. Bar, 10 μm. The co-localization index of MMP14 with indicated markers (EEA1, Rab7 and LAMP1) was determined by the measurement of overlapped signaling (yellow fluorescence) over total MMP14 signal. Data are shown as box plots together with individual data points, and whiskers indicate minimum to maximum (n=6 independent experiments). **, P < 0.01, ***, P < 0.001 as determined by unpaired two-tailed t-test. **(C-D)** Time-course analysis of MMP14 protein levels after cycloheximide (CHX) treatment. Ctrl or ATP6AP2-KD MC3T3 cell lines were treated with 50 µg/ml CHX for the indicated time. ATP6AP2 and MMP14 protein levels were analyzed by Western blotting. Representative blots are shown in C, and quantification analysis (mean ± SD from three separate experiments) is presented in D. P values obtained by two-way ANOVA followed by Bonferroni post hoc test. *, P < 0.05. **, P < 0.01.

